# Drought tolerance of *Aedes aegypti* mosquito eggs is influenced by adaptation to local climate conditions and associations with humans

**DOI:** 10.64898/2026.01.15.699549

**Authors:** Souvik Chakraborty, Emily Zigmond, Kaylee Levan, Jasmine Kennedy, Akshita Vaka, Thomas Arya, Sher Shah, Melissa Uhran, Angelic Harris, Massamba Sylla, Jewelna Akorli, Sampson Otoo, Noah Rose, Christopher J. Holmes, Yanyu Xiao, Joshua B. Benoit

## Abstract

Urbanization is intensifying human interactions with mosquitoes, exacerbating public health challenges. Densely populated areas provide ideal conditions for container-dwelling mosquitoes, with increased host availability and the presence of artificial breeding sites. These anthropophilic mosquitoes often exhibit distinct ecological adaptations compared to their rural counterparts. Since mosquito eggs are immobile and remain at the site of oviposition, they provide a valuable lens for assessing how urbanization, climate-driven shifts in temperature, and drought affect mosquito reproductive success. This study examined *Ae. aegypti* egg viability under varying temperature and dry conditions over five months, focusing on lineages with distinct ancestries from West African populations. Mosquitoes collected from urban habitats with a high human preference demonstrated higher egg survival under prolonged arid conditions. Analysis of climatic factors revealed that dry season temperature and precipitation during wet periods are significant predictors of egg drought tolerance. Modeling future climate scenarios based on input from our egg viability results suggests a projected shift and expansion in the seasonal survival window for *Ae. aegypti* by the end of the century. This study highlights the importance of understanding environmental constraints on the drought tolerance of mosquito eggs to predict and mitigate future mosquito outbreaks.

## Introduction

Urbanization, growing human population, and subsequent anthropogenic warming are significant ecological drivers reshaping interactions with arthropods that vector disease, posing critical public health challenges (Kache et al., 2022; Wilke, Benelli, et al., 2021). *Aedes aegypti* is a vector for diseases such as Zika, chikungunya, dengue fever, and yellow fever (Sukhralia et al., 2019; Weaver et al., 2018). Inadequate sanitation, poor water management, long-term unplanned construction-sites in urban areas increase habitats conducive to *Ae. aegypti* breeding, amplifying risk associated with mosquito-borne pathogens (de Souza & Weaver, 2024; Gibb et al., 2023). Simultaneously, the geographic range of *Ae. aegypti* is expanding, introducing disease transmission risks into previously inhospitable regions (de Souza & Weaver, 2024). Central to this range expansion is the resilience of *Ae. aegypti* eggs to environmental stressors, enabling persistence during unfavorable conditions for months until rainwater triggers hatching (Juliano et al., 2002; Rezende et al., 2008; Sota & Mogi, 1992). These patterns in *Ae. aegypti* reflect broader evolutionary trends seen across insects, where egg size, shape, and morphology evolve primarily in response to ecological pressures such as habitat type and environmental dryness, with oviposition site playing a key role in shaping these traits (Church et al., 2019; Sota & Mogi, 1992). Although *Aedes* eggs persist, this stage is vulnerable, as eggs are typically immobile and restricted to the location of deposition (Chu et al., 2022), unless transported via human-mediated trade while still attached to surfaces (Powell et al., 2018). Temperature and humidity are critical environmental factors that influence egg success, with each exerting pressure on egg viability (Ajayi et al., 2024; Bowler & Terblanche, 2008; Fischer et al., 2025; Guarneri et al., 2002; Sota & Mogi, 1992). Our recent work found that urban *Ae. aegypti* populations exhibit increased thermal tolerance, with higher viability near their upper thermal limits than rural-origin populations, suggesting adaptation to urbanization (Chakraborty, Zigmond, et al., 2025). In general, *Ae. aegypti* eggs exhibit remarkable desiccation resistance (Faull et al., 2016; Juliano et al., 2002; Meola, 1964; Sota & Mogi, 1992; Trpis, 1972). Comparative work across *Aedes* species shows that desiccation survival is shaped by habitat type, with forest-associated species generally exhibiting reduced egg survival compared to urban-adapted species such as *Ae. aegypti aegypti* (Sota & Mogi, 1992). These insights on mosquito egg desiccation have been gained from limited stress conditions without exposure to varying thermal conditions (Faull & Williams, 2015; Fischer et al., 2025; Meola, 1964). This gap underscores the need to assess egg survivorship across a broader range of dehydration regimes, particularly for a species adapted to contrasting habitats.

*Ae. aegypti* is native to Africa and originated there as a forest-dwelling species, feeding on various animals. A significant evolutionary shift occurred in the West African Sahel, where a subset of the population specialized in anthropophilic biting and domestic habitats, driving the divergence into two distinct subspecies. The anthropophilic, domestic *Ae. aegypti aegypti* (*‘Aaa’*), predominantly utilizes artificial containers for oviposition, and the generalist *Ae. aegypti formosus* (*‘Aaf’*), the more versatile counterpart, favors natural egg-laying sites (McBride et al., 2014; Rose et al., 2020). Artificial containers often create warmer and more humid microclimates than natural environments. Although these conditions may initially enhance egg viability, they simultaneously elevate the risk of desiccation as these artificial containers are prone to rapid drying. The contrasting oviposition behaviors of *Ae. aegypti* subtypes provide a unique model for studying the interplay between egg-hatching biology, ecological adaptation, and the broader implications of global climate change (Chakraborty, Zigmond, et al., 2025; Fischer et al., 2025). Understanding intraspecific variation in egg desiccation tolerance is essential for elucidating the human specialization continuum in *Ae. aegypti*, their global spread, and species survival under climate change scenarios (Juliano & Philip Lounibos, 2005; Martín et al., 2025). Notably, in the dry parts of African Sahel, smaller towns predominantly harbor anthropophilic ‘*Aaa’* rather than *‘Aaf’* (Rose et al., 2020), though this pattern may not generalize across regions. Similarly, populations from a high human density area, ‘Ngoye,’ represent human-specialist mosquitoes, even though it is more of a rural-town where natural tree-hole habitats are scarce, which likely forced a dependence on humans for survival (Rose et al., 2020, 2023). Increased human populations and urban activities contribute to the formation of heat islands, (Hsu et al., 2021; Tuholske et al., 2021) where warmer, more humid microhabitats can serve as refuges during extreme dry periods, enabling rapid population expansion following rainfall. Thus, anticipating egg-level traits, such as desiccation tolerance, becomes central to predicting mosquito population persistence and outbreak potential under future climate extremes. This becomes especially important in regions away from the tropics, as climate warming expands poleward, where previously cooler regions are experiencing conditions more conducive to mosquito persistence (de Souza & Weaver, 2024).

This study investigates the hatching of *Ae. aegypti* eggs under different temperature and humidity conditions over five months, focusing on urban- and rural-origin populations from West Africa. We evaluate the relationship between egg-drought tolerance and ecological factors, including bioclimatic variables, human population density, mosquito ancestry, and their host preference. With our experimental data we bridged climate data and explored future mosquito population dynamics throughout a calendar year over the end of this century. We modeled location-specific mosquito trajectories under various Shared Socioeconomic Pathway (SSP) scenarios across African study sites and extended our analysis to three locations each in India and the USA. These sites were selected based on *Ae. aegypti* prevalence, human population density contrasts, recent outbreak history, climate projections, and study objectives (H. C. et al., 2024; Kakarla et al., 2020; Varamballi et al., 2024; Wilke, Vasquez, et al., 2021; Yadav et al., 2019). We developed a novel predictive pipeline that integrates historical and projected future climate data to estimate yearly trends in mosquito viability and adult survival under climate change.

## Materials and methods

### Mosquito collection, maintenance, and experimental protocol

Seven *Aedes aegypti* mosquito colonies were established from several locations across Sub-Saharan West Africa (Rose et al., 2020), with urban and rural classifications determined primarily by human population density (Supplementary Table 1; (Chakraborty, Zigmond, et al., 2025)- Table 1). Mosquito populations were collected between 2017 and 2018, and experiments were conducted between 2021 and 2023. The colonies were maintained in a climate-controlled facility at a temperature of 27±2 °C, with a relative humidity (RH) of 65±4%, and a 16:8 hour light-dark cycle. The mosquito larvae were reared in distilled water with finely pulverized fish food (Tetramin, Goldfish Flakes) and yeast extract (Difco, BD 210929) (Chakraborty, Zigmond, et al., 2025). The pupae were placed in cups, and the adult mosquitoes were housed in 60 cm × 60 cm × 60 cm mesh cages with access to water and 10% sucrose solution through soaked cotton wicks. After reaching an age of 10-14 days, female mosquitoes were fed on a human host (University of Cincinnati IRB 2021-0971), and the eggs were collected on brown paper towels. The eggs were kept in the climate-controlled colony facility for two weeks to facilitate embryonation before experiments (Maïga et al., 2017). A batch of eggs (50-70 eggs) was placed in 8 milliliter size plastic vials (n= 10 vials/treatment) with punched lids, and the plastic vials were housed inside desiccators for the designated treatment time. These desiccators were placed in temperature-controlled incubators to maintain precise conditions and to verify and monitor the temperature and humidity inside the desiccators; data loggers (ONSET HOBO) were placed within. Nine temperature-humidity combinations were designed with three temperature levels (23, 29, and 35 [±1] °C) alongside three humidity levels (33, 55, and 75 [±2] % RH) (Supplementary Figure 1). Eggs were assessed after 1, 4, 8, 12, 16, and 20 weeks.

**Table 1:** Summary of mixed-effects model comparisons assessing mosquito egg drought tolerance. Models include various combinations of predictors, such as genetic ancestry, host preference, population density, human specialization index, and bioclimatic variables. A random intercept for mosquito line is included in all models to account for repeated measures. Model performance is evaluated using Akaike Information Criterion (AIC) and likelihood ratio tests (χ² statistics and associated *p*-values) comparing each model against a reference to determine relative fit and statistical significance.

| Model | Comparison | AIC | Chisq | Df | PrChisq |
| --- | --- | --- | --- | --- | --- |
| null_model(Ho) |  | -23.39137666 | 0 | 0 |  |
| Imm_pref (Hp) | Ho VS Hp | -21.32533431 | 4.671673986 | 1 | 0.030664025 |
| pref_interaction (Hp') | Ho VS Hp' | -15.69658212 | 9.616183701 | 2 | 0.008163422 |
| Imm_ancestry (Ha) | Ho VS Ha | -24.99322924 | 5.907544309 | 1 | 0.015076174 |
| ancestry_interaction (Ha') | Ho VS Ha' | -22.17284283 | 11.20296425 | 2 | 0.003692387 |
| model_(weighted)_Preference_Ancestry (H_a_p) | Ho VS H_a_p | -11.612834 | 11.82486209 | 4 | 0.018702283 |
| Imm_density (Hd) | Ho VS Hd | -19.91678274 | 6.080833771 | 1 | 0.013665626 |
| density_interaction (Hd') | Ho VS Hd' | -10.83900583 | 10.25340978 | 2 | 0.005936088 |
| model_density_p_a_weighted (H_hss) | Ho VS H_hss | -7.167493213 | 11.56013451 | 4 | 0.02094029 |
| model_(weighted)_Preference_Ancestry VS.<br>model_density_p_a_weighted | H_a_p VS H_hss | ΔAIC =3.748 | 7.7474 | 2 | 0.02078 |
| bio1 (H_bio1) | Ho VS H_bio1 | -19.71390607 | 4.331160965 | 1 | 0.037420714 |
| bio1_interaction (H_bio1') | Ho VS H_bio1' | -10.13944319 | 6.337239525 | 2 | 0.042061613 |
| bio3 (H_bio3) | Ho VS H_bio3 | -19.68694598 | 7.634311344 | 1 | 0.005726825 |
| bio3_interaction (H_bio3') | Ho VS H_bio3' | -8.447093111 | 11.30649408 | 2 | 0.003506114 |
| bio4_main (H_bio4) | Ho VS H_bio4 | -12.59699131 | 5.306683157 | 1 | 0.021243764 |
| bio4_interaction (H_bio4') | Ho VS H_bio4' | 4.822932798 | 7.534391612 | 2 | 0.023116796 |
| bio9 (H_bio9) | Ho VS H_bio9 | -20.67647717 | 5.431763734 | 1 | 0.019773699 |
| bio9_interaction (H_bio9') | Ho VS H_bio9' | -14.02976179 | 10.7684086 | 2 | 0.00458849 |
| bio10 (H_bio10) | Ho VS H_bio10 | -19.85366823 | 6.010456073 | 1 | 0.014221351 |
| bio10_interaction (H_bio10') | Ho VS H_bio10' | -9.680518675 | 8.968033405 | 2 | 0.011287982 |
| bio11 (H_bio11) | Ho VS H_bio11 | -20.70831156 | 5.062734634 | 1 | 0.024445636 |
| bio11_interaction (H_bio11') | Ho VS H_bio11' | -11.71091017 | 7.377615795 | 2 | 0.025001789 |
| bio13 (H_bio13) | Ho VS H_bio13 | -11.65540622 | 4.382816655 | 1 | 0.036302959 |
| bio13_interaction (H_bio13') | Ho VS H_bio13' | 6.824255355 | 5.507444593 | 2 | 0.046369035 |
| bio16 (H_bio16) | Ho VS H_bio16 | -9.787749763 | 4.82242129 | 1 | 0.028091857 |
| bio16_interaction (H_bio16') | Ho VS H_bio16' | 10.63285548 | 6.328789281 | 2 | 0.042239704 |
| bio18 (H_bio18) | Ho VS H_bio18 | -9.091213585 | 3.390572851 | 1 | 0.046557016 |
| bio18_interaction (H_bio18') | Ho VS H_bio18' | 9.858636686 | 5.785159024 | 2 | 0.045543304 |
| model_climate (H_bio9') | Ho VS H_bio9' | -14.02976179 | 10.7684086 | 2 | 0.00458849 |
| model_density_p_a_weighted_bio9 (H_hss_climate) | Ho VS H_hss_climate | 8.21342 | 12.101 | 6 | 0.05975 |
| model_density_p_a_weighted VS model_climate | H_hss VS H_bio9' | ΔAIC =3.208 | 0.7917 | 2 | 0.6731 |
| model_density_p_a_weighted VS | H_hss VS H_hss_climate | ΔAIC =3.459 | 0.5411 | 2 | 0.094 |

At a specific time, eggs were removed from incubators, and DI water with fish food was pipetted into the vials to prompt hatching. Following larval emergence, around 40-48 hours later, 80% ethanol was pipetted into the vials for storage and future counting. During the egg-counting process, we counted the total number of hatched and unhatched eggs. Hatched eggs were identified from unhatched eggs by a clean cut/break of the egg cap, indicating the emergence of the larvae, where unhatched eggs remained intact, showing no evidence of larval emergence. The egg survival rate was assessed using two complementary methods. First, we calculated the hatching ratio, defined as the number of hatched eggs divided by the total number of eggs, providing a quantitative measure of the proportion of eggs that hatched. Second, we employed a binary outcome framework, using cbind(success, failure), to classify outcomes as hatching or non-hatching. Both approaches yielded consistent results, and we used the first method to determine egg drought tolerance (Chakraborty, Zigmond, et al., 2025).

### Egg drought tolerance and temporal analysis of key predictors for egg drought tolerance

Saturation deficit (SD), a measure of air dryness that incorporates saturation vapor pressure (determined by temperature) and actual vapor pressure (percent relative humidity, at the given temperature), was used to investigate predictors of egg drought tolerance (Holmes & Benoit, 2019; Linde et al., 1990). To estimate the saturation deficit, we used the Tetens Equation or saturation vapor pressure of water (P_sat_) to calculate the vapor pressure deficit (VPD), resulting in kilopascals, and subsequently converted to millibars (mbar). Each combination of temperature and RH produced nine distinct saturation deficit (SD) conditions, ranging from 7 to 37 mbar (Supplementary Figure 1). To assess population-specific drought resilience, deviations in egg hatching were analyzed for each time-treatment combination relative to the overall hatching rates across all populations. Egg hatching was analyzed at different thresholds, with lethal times for 50% and 75% viability. Temporal trends were explored by categorizing time points into “early” (1, 4, 8 weeks) and “late” (12, 16, 20 weeks) periods. This timing captures ecologically meaningful differences, as we noted that egg drought tolerance in *Ae. aegypti* appears to diminish significantly onwards 12 weeks (Juliano et al., 2002). Non-parametric Wilcoxon rank-sum tests were performed to compare egg-hatching responses between urban and rural populations at each time-treatment condition to assess habitat-based differences in drought tolerance. This analysis provided foundational insights into temporal trends and variations in egg-hatching responses among mosquito populations originating from areas with contrasting human population density.

Following the initial identification of significant differences in egg survival across time points and treatments between urban- and rural-origin mosquito populations, we next examined the relationship between egg hatching and SD for each population at each time point. Using a simple linear model (Egg hatching = m × SD + c), we analyzed experimental hatching outcomes across nine SD conditions at each ‘week’ level. This modeling framework captured the temporal dynamics of egg drought tolerance, reflected in the regression coefficients (‘m’), and served as the foundation for projecting experimental hatching patterns onto future climatic scenarios to forecast mosquito population responses to environmental change. While exploring egg drought tolerance, two approaches were considered here: 1) “m×SD+c” and 2) “m”. Consistent results were obtained by both approaches. Ultimately, we used model (2) for further analysis, representing each population with a single regression factor, i.e., the slope derived from the linear model (Chakraborty, Zigmond, et al., 2025). The regression coefficient ‘m’ serves as a standalone proxy for desiccation vulnerability, where the magnitude of the slope represents the intrinsic sensitivity of egg viability to atmospheric moisture across each population and experimental condition.

Lastly, to investigate the biological and environmental determinants of egg drought tolerance, we employed a linear mixed model (LMM) approach. Specifically, this analysis aimed to elucidate the adaptive divergence and ecological specialization of the two *Ae. aegypti* subspecies, ‘*Aaa*’ and ‘*Aaf*’ under climatic pressure (McBride et al., 2014; Rose et al., 2020). These findings were interpreted within the framework of “human specialization syndrome” (HSS) (Chakraborty, Zigmond, et al., 2025), which describes a suite of traits, including genetic background and behavioral shifts driven by prolonged exposure to high human population densities, enhancing specialization on human hosts (Rose et al., 2020). Moreover, eleven temperature-related and eight precipitation-related bioclimatic variables (Fick & Hijmans, 2017) were modeled to test their temporal impact on egg drought tolerance over five months using the LMM approach, assessing each variable individually and in combination. LMMs handle within-subject correlations more accurately by incorporating random effects (e.g., ‘line’), offering a more precise account of egg drought tolerance over time (Arnau et al., 2012; Cnaan et al., 1997). Significant interactions between each predictor and time (‘week’) suggest that the strength and nature of these associations vary throughout the experiment. Correlation heatmaps visualized the temporal associations between each bioclimatic variable and egg drought tolerance, with statistical significance assessed for each week. We used the Akaike information criterion (AIC) and Bayesian information criterion (BIC) to select the best-fitting model. All statistical analyses were conducted in R v. 4.2.3 using the “lm” function for linear models and the “lmer” function for LMMs (Kuznetsova et al., 2017). Figures were also produced using R v. 4.2.3 and edited with Inkscape 1.3.

### Climate and mosquito population projections

To explore the influence of temperature-dependent traits on mosquito population dynamics, we first quantified the effects of temperature on survival, development, oviposition timing, and fecundity across life stages (Supplementary method). The temperature-dependent life stage-specific survival rates were calibrated based on the datasets from earlier studies in which thermotolerance (Chakraborty, Zigmond, et al., 2025; Couret & Benedict, 2014) and mosquito population dynamics were experimentally measured (Simoy et al., 2015; Yang et al., 2009). We further converted these temperature-dependent metrics into time-dependent functionals by integrating historical climate records with monthly temperature projections from the Intergovernmental Panel on Climate Change (IPCC) Sixth Assessment Report for each mosquito collection and case study site (*Climate Change Knowledge Portal*, 2025). Then these calibrated functionals were incorporated into an age-structured population model, which allows us to project future mosquito populations for various locations. Simulations were initialized with 20 individuals per stage (egg, larva, adult) to understand the pattern of mosquito survival and emergence for each study site. The model divides the population into five developmental groups (Supplementary method), including eggs, larvae, newly emerged adults in their first gonotrophic cycle before oviposition, adults in their resting period and subsequent cycles before oviposition, and adults in oviposition during their subsequent cycles (Holmes et al., 2025).

### Relative humidity as a critical dimension to evaluate survival of eggs

While temperature is often considered the primary driver of mosquito population dynamics, RH is an essential yet frequently overlooked variable in thermal biology and mosquito-borne disease transmission (Brown et al., 2023; Lega et al., 2017). To refine our understanding of mosquito survival, we incorporated RH predictions into our analysis (Supplementary method). RH estimates were derived for historical datasets from IPCC; however, given the uncertainties in RH estimates across different study locations, we employed a formula to generate future model-predicted values, as, RH_h_/T_h_ = (RH_f_ - RH_p_)/ (T_f_ - T_p_); where T = temperature, and RH = relative humidity and the subscript index h = historical conditions, f = future conditions, p = model-predicted values. A truncated Fourier series (sine-cosine) function was fitted to capture the seasonal variation in historical temperature and RH across calendar months using the “nls” function in R v. 4.2.3 (Baty et al., 2015; Estrada-Peña et al., 2014) to generate the model-predicted climatic elements. The egg-hatching patterns were then visualized against both IPCC-sourced climatic data and our cross-validated predictions; both patterns closely mirrored each other, and we ensured consistency throughout the experiment using our own validated RH dataset, generated through model prediction values. As in previous analyses, SD was computed from temperature and RH to quantify air dryness, allowing us to visualize egg-hatching patterns using the fitted values from experimental data. To ensure biological relevance in modeling egg hatching and mosquito population survival, we applied several entomological transformations. Predictions for mosquito egg hatching incorporated climatic conditions from prior months, recognizing that *Ae. aegypti* eggs remain viable for extended periods. Early egg-hatching predictions were based on climatic conditions from the preceding two months, while late-hatching predictions incorporated data from the prior four months. This categorization captures the ecological dynamics whereby rainfall can initiate immediate hatching or be retained in natural and artificial containers, maintaining microhabitats that support delayed egg hatching. Additionally, a lower temperature threshold for biological survival was set at 10°C, below which all mosquito life stages were assumed to perish. Egg hatching success was estimated using best-fit models derived from our experimental data on egg drought tolerance. Larval and adult survival probabilities were calculated using a previous study (Chakraborty, Zigmond, et al., 2025). These stage-specific probabilities were multiplied to compute the overall survival probability and to project monthly occurrence patterns over the coming decades, capturing the cumulative impact of climate on *Ae. aegypti* under various SSP scenarios.

## Results

### Egg survival of mosquito eggs

Urban-rural differences in egg hatching were significant at the two highest saturation deficit (SD) levels across all time points, with additional differences only appearing at weeks 16 and 20 (Supplementary Figure 2). Egg-hatching patterns across all populations show a high degree of overlap at lower SDs (Figure 1a), with a divergence between urban and rural populations exceeding 25 millibars (mbar), mostly 12 weeks onward (Figure 1b; Supplementary Figure 3). When SD remains below 20 mbar, egg viability showed relative stability, declining only by 25-30% over 20 weeks (Supplementary Figure 4). Maximum egg hatching occurred at early time points (1, 4, 8 weeks) under experimental wet conditions (SD < 15 mbar). After 20 weeks, mean egg hatching at the highest SD (37.66 mbar) was nearly 2-fold greater in urban populations (0.2 ± 0.08) than in rural populations (0.07 ± 0.04; Figure 1b). At this timepoint, urban-origin ‘Ngoye’ showed the highest hatching (0.22 ± 0.05), while ‘PK10’ (rural-origin) showed the lowest (0.04 ± 0.02) (Supplementary Table 2). Although egg hatching never dropped below 50% during the first two months under any experimental conditions, the urban-rural dichotomy in egg hatching remained consistent at the onset of the third month (linear mixed-effects model, *p* < 2 × 10⁻^16^) (Supplementary Figure 4). These results highlight a critical threshold after the third month, beyond which egg viability declines sharply, reinforcing the importance of SD in driving adaptive differences between urban and rural lineages. Egg-hatching thresholds (LT_50_ and LT_75_) were calculated across SD conditions; however, egg hatching at low (<15 mbar) and moderate (15–25 mbar) SDs did not fall below 75% (Supplementary Figure 3), identifying the highest three stress levels (25.3, 26.83, and 37.66 mbar) as the most critical to egg viability (linear mixed-effects model, *p* < 2 × 10⁻^16^) (Supplementary Table 3). Among urban populations, Ngoye demonstrated the highest drought resistance, with an LT_50_ of 11.71 weeks at the driest SD condition (37.66 mbar). In contrast, rural populations like PK10 (9.99 weeks) and Kedougou (9.64 weeks) displayed the lowest LT_50_ values under the same SD condition, reflecting reduced drought tolerance (Supplementary Table 3; Supplementary Figure 5).

**Figure 1:**
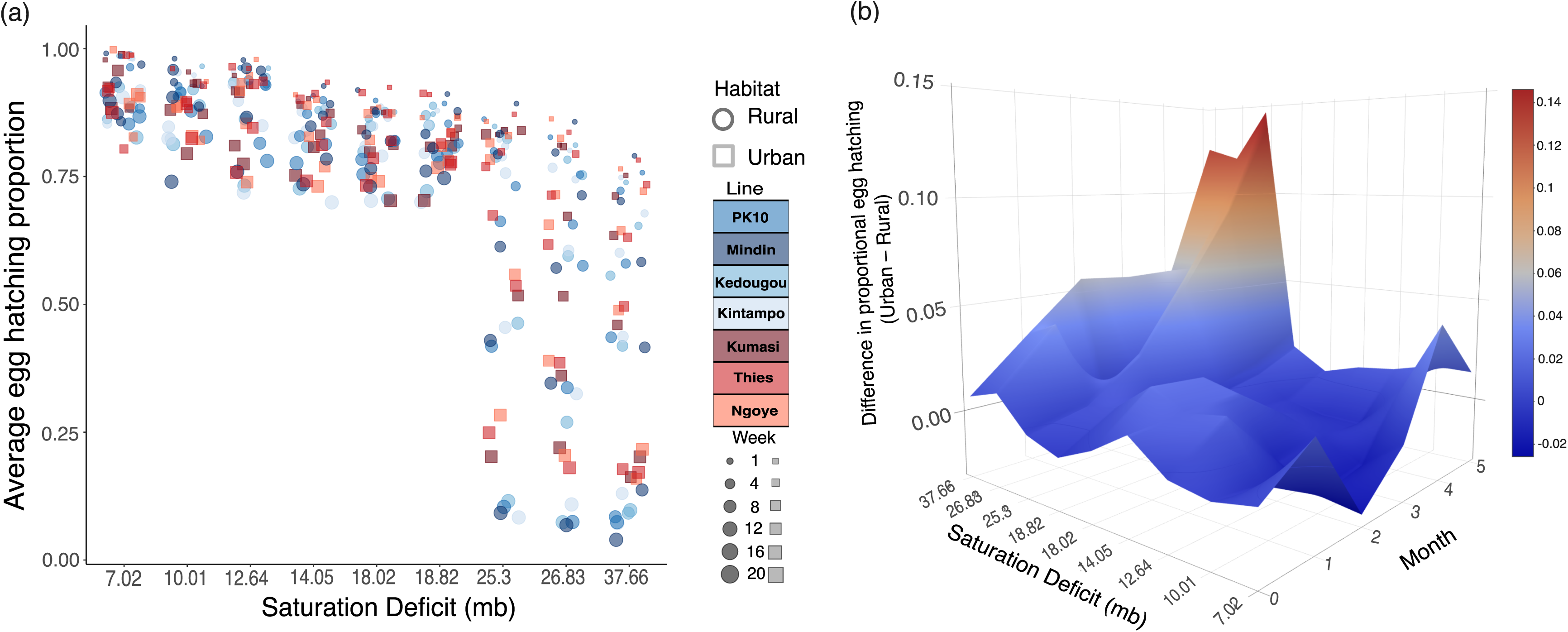
Survivorship of *Aedes aegypti* eggs under saturation deficit (SD) stress (mbar). (a) Average egg hatching across *Ae. aegypti* populations following exposure to SD conditions for up to 20 weeks. Dot color and shape indicate habitat type (blue shade and circle = rural, red shade and square = urban) and dot size reflects the number of weeks under stress. (b) Habitat-specific differences in egg hatching (urban-rural) after SD stress for up to 20 weeks. Statistical analyses were performed using generalized linear modeling, when necessary.

### Evaluating temporal influences of egg drought tolerance in *Aedes aegypti*

Rural populations exhibited a steeper decline in egg drought tolerance over-time compared to urban populations (linear mixed-effects model, *p* < 0.013), indicating heightened sensitivity to prolonged arid conditions (Figure 2a). While early differences were evident, by weeks 16 and 20, urban populations consistently maintained higher egg drought tolerance. Building on these temporal patterns, we further analyzed the relative contributions of the human specialization syndrome (HSS) and key bioclimatic variables to egg drought tolerance (Table 1), with most of the associations occurring during later stages in weeks 12, 16, and 20 (Figure 2b). All three HSS components, i.e., (1) ancestry, related to (2) host preference and (3) local human population density were significantly associated with egg drought tolerance (Table 1). These variables functioned both as individual predictors and as temporal (week-level) interactions, highlighting that their influence is not static but evolves as drought conditions persist. Given the known association between ancestral origin and host preference (Chakraborty, Zigmond, et al., 2025; Rose et al., 2020), we generated a weighted composite of these two traits, which showed a significant association with egg drought tolerance (Table 1; LRT; χ² = 11.825; *p* = 0.02). Subsequently, adding human population density to the model further improved the fit viz., ΔAIC = -5.449 (Table 1; LRT; χ² = 9.45 *p =* 0.009), suggesting that density contributes additional explanatory power beyond ancestry and preference.

**Figure 2:**
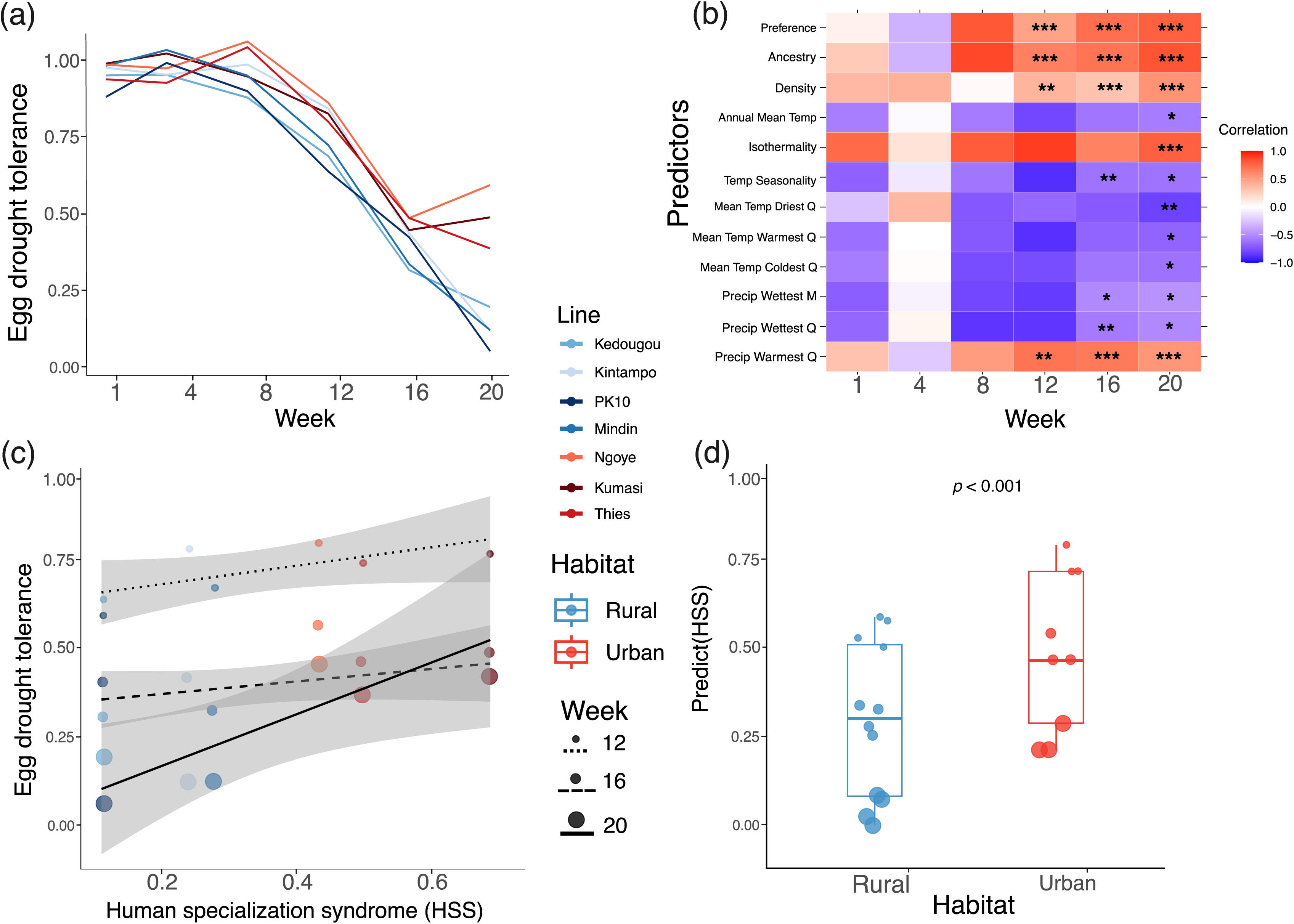
Temporal dynamics of biological drivers influencing egg drought tolerance in Aedes aegypti. (a) Population-specific trends in temporal egg drought tolerance across consecutive weeks. Drought tolerance was statistically quantified using proportional analysis and cbind(success, failure) independent methods, each yielding consistent patterns. (b) Key predictors influencing egg drought tolerance over time, as identified through a linear mixed-effects modeling framework (Table 1; Supplementary Table 1). Tile color indicates the direction and strength of temporal correlations with drought tolerance. Significance is denoted as follows: *** *p* < 0.001, ** *p* < 0.01, * *p* < 0.05. (c) The “human specialization syndrome” (HSS) provided the best fit for explaining time-dependent variation in egg drought tolerance, with distinct temporal patterns across HSS classifications (Table 1). (d) Predicting through the lens of human specialization, a clear divergence in egg viability emerges between urban and rural populations (LRT; *p* < 0.001), revealing habitat-specific adaptive responses to desiccation stress.

Among 11 temperature-related bioclimatic variables, annual mean temperature (bio1), isothermality (bio3), temperature seasonality (bio4), and mean temperatures of the driest (bio9), warmest (bio10), and coldest (bio11) quarters emerged as significant predictors of egg drought tolerance, both individually and over time (Table 1). Similar to the HSS, these climatic effects were largely associated with egg drought tolerance at the later experimental stages (Figure 2b). Notably, the mean temperature of the driest quarters, bio9, emerged as the strongest predictor among temperature variables (LRT; χ² = 5.43; *p* = 0.02). For eight precipitation-related bioclimatic variables, significant associations were found with precipitation during the wettest month (bio13), wettest quarter (bio16), and warmest quarter (bio18), with bio13, and bio16 showing the most consistent and statistically robust association (Table 1), supported by AIC, BIC, and log-likelihood comparisons. Among all the 19 bioclimatic variables tested, bio9 (mean temperature of driest quarter) showed only a significant association with HSS, regarding egg drought tolerance (LRT; χ² = 11.98; *p* = 0.03).

However, the HSS model incorporating weighted ancestry and preference, along with human population density, performed best overall (Table 1; Figure 2c), slightly outperforming a model based solely on climate variables (χ² = 0.7917; *p* = 0.67) or the combined HSS-climate index (χ² = 0.5411; *p* = 0.09). Results from a linear mixed-effects model indicated that habitat-specific egg drought tolerance differs significantly with HSS (Figure 2d); after accounting for temporal declines (weeks), urban habitats exhibited higher predicted HSS values compared to rural habitats (*p* < 0.001). Our results consistently show that human-associated traits each contribute uniquely to egg desiccation resistance, functioning as complementary and non-redundant drivers of drought adaptation. These effects operate in parallel with key local temperature and precipitation variables, particularly in dry- and warm-season dynamics, and are likely to play a greater role as anthropogenic warming accelerates.

### Predictive modeling of temperature-driven mosquito population dynamics

Temperature-dependent ovipositional delay, egg output, and developmental duration were incorporated into life stage-specific survival models to estimate potential mosquito population growth. Urban and rural habitats of African study locations exhibited similar seasonal temperature patterns (Figure 3a), but life-history traits diverged notably. Urban mosquitoes showed shorter oviposition intervals (Figure 3b), longer egg and larval development times (Figure 3c), and higher adult survival rates (Figure 3d). Egg production was slightly higher in rural populations during peak reproductive months (Figure 3e). However, mosquito population sizes were substantially higher in urban areas (Figure 3f), underscoring the heightened vector potential in urbanized environments. Starting with an initial cohort of 20 individuals per life stage (eggs, larvae, and adults), our predictive modeling demonstrates how temperature influences mosquito population dynamics across geographies (Figure 3f and 4). Simulations revealed that adult mosquito populations (ranging from 100 to 100 millions) could re-establish and grow rapidly within one to two years from the initial cohort (Figure 4). Although these simulations are area-dependent, as in Ohio, a temperate zone, cold winters severely limit egg and adult survival. The largest projected increases occurred in tropical and near-tropical regions (Figure 4a,b,c), where warm and humid conditions accelerate life cycle turnover and enhance reproductive success. Notably, Indian locations, i.e., Delhi, Karnataka, and Rajasthan, indicated substantial population growth beginning in the second year of simulation, where population surge crossed millions in number. In contrast, mosquito populations in U.S. locations, Ohio, Arizona, and Florida, showed earlier emergence, typically within the first year of simulation. However, the overall magnitude of expansion was markedly lower than in Indian regions. Additionally, smaller initial population sizes further constrained recovery potential. Although, in Florida, the mosquito population displayed a more favorable pattern for growth, with emergence of adults observed within the first year from the mere founder population. The variety in emergence timing of adults and population recovery may reflect broader regional disparities in geography, urbanization, and anthropogenic pressures. Although issues such as pollution, land use, and urban heat islands certainly influence vector dynamics, these effects were not explicitly incorporated in the current temperature-driven simulations.

**Figure 3:**
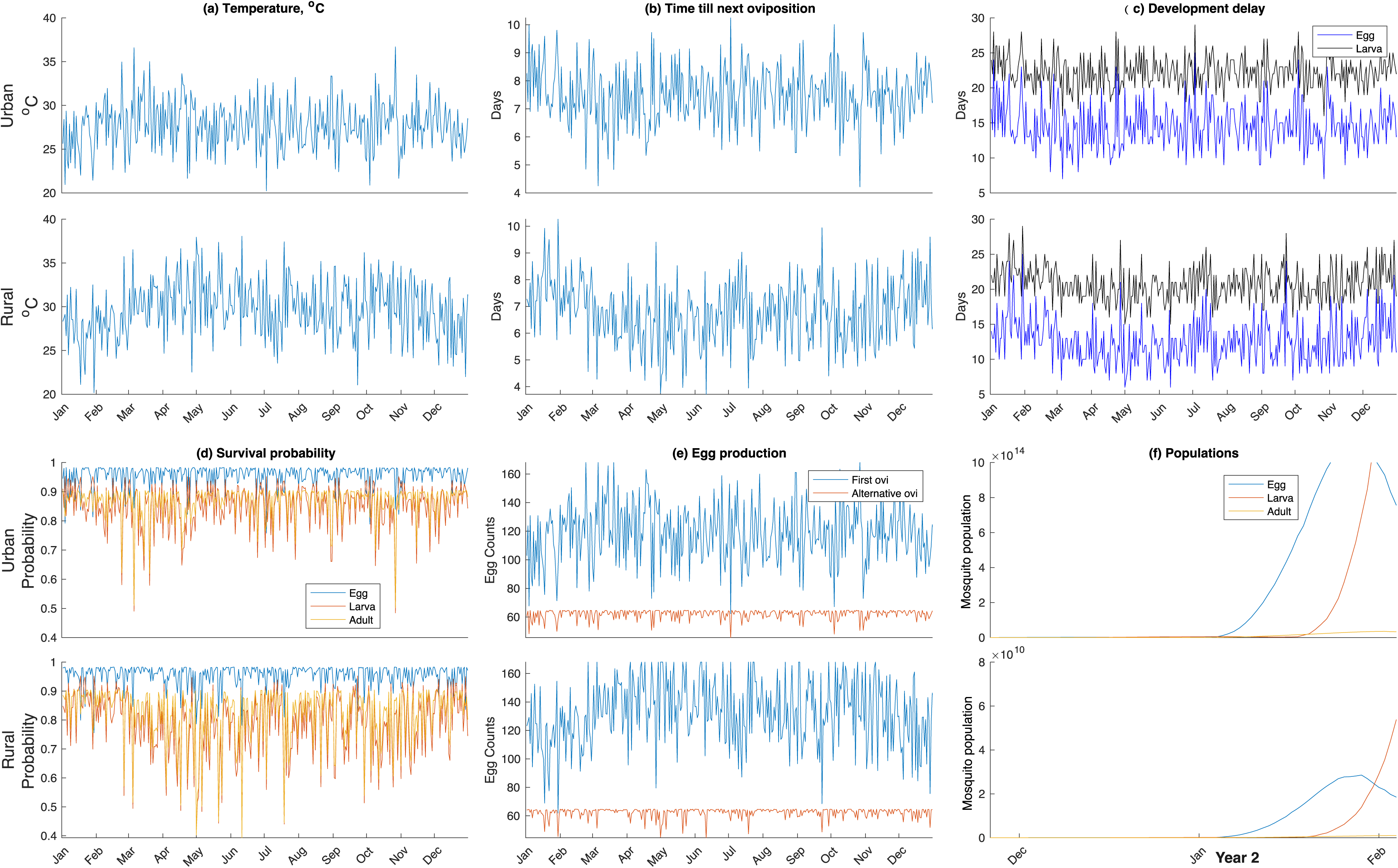
Predictive, temperature-based modeling of *Aedes aegypti* population dynamics across urban and rural habitats of African study sites. Simulations for urban and rural lineages by integrating seasonal temperature profiles with empirically derived biological trait transformations (Supplementary modeling). The model outputs for urban and rural lineages include: (a) seasonal temperature trends, (b) oviposition intervals, (c) development time from egg to larva and larva to adult, (d) stage-specific survival rates, (e) monthly egg production during the first gonotrophic cycle and subsequent cycles, and (f) projected population expansion from an initial cohort of 20 mosquitoes. Temperature-dependent traits such as thermotolerance, reproductive output, developmental delays, and survival rates were informed by experimental data (Chakraborty, Zigmond, et al., 2025; Couret & Benedict, 2014; Simoy et al., 2015; Yang et al., 2009).

**Figure 4:**
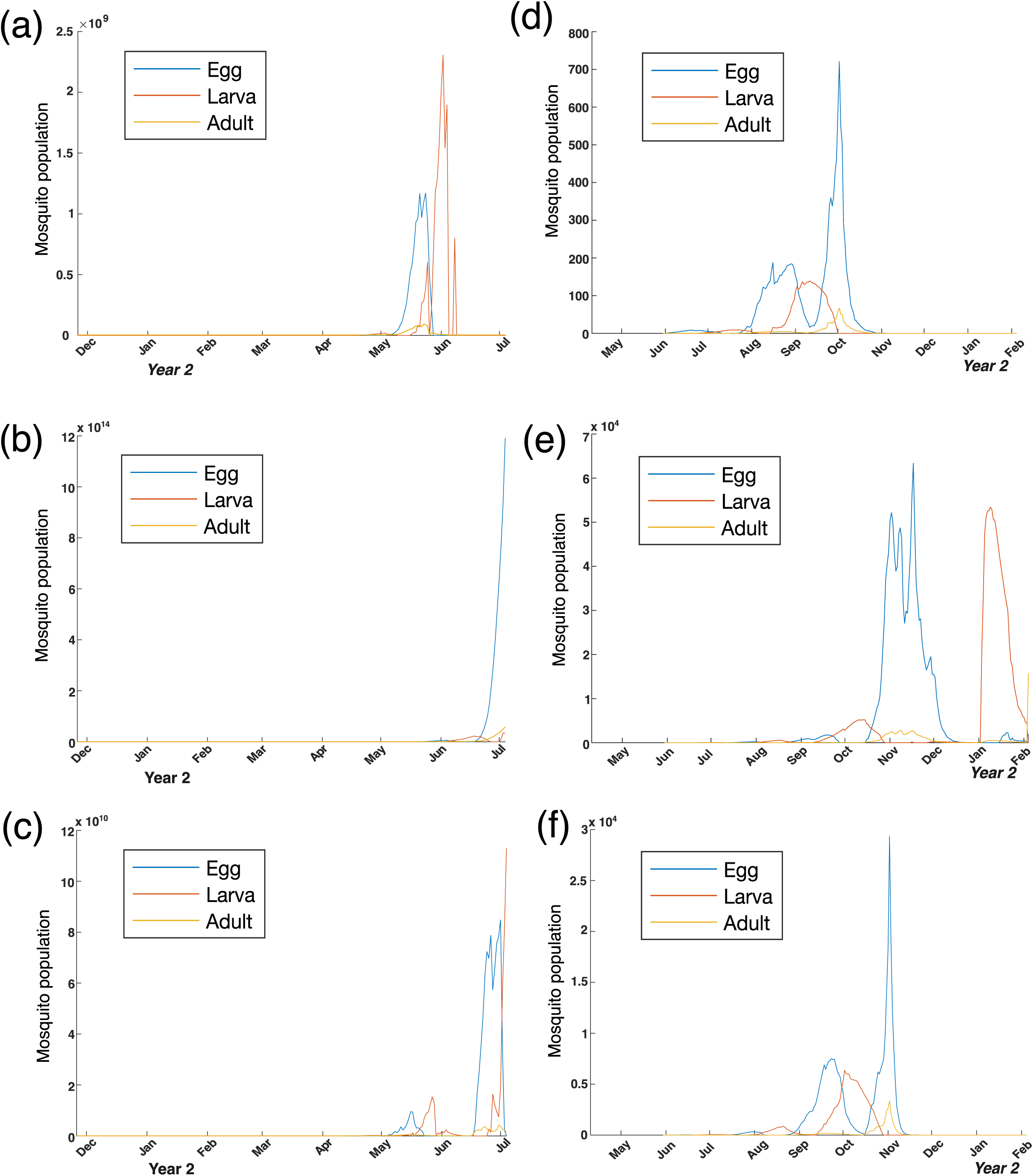
Comparative modeling of *Aedes aegypti* population outbreak timing across six case study locations. Predictive simulations were conducted for three Indian states, (a) Delhi, (b) Karnataka, (c) Rajasthan and three U.S. states, (d) Ohio, (e) Florida, (f) Arizona to evaluate potential mosquito population surges under site-specific seasonal temperature regimes. Model outputs reflect projected population dynamics from an initial cohort of 20 mosquitoes. The timing of peak population emergence differs by panel and is projected on the period of most significant outbreak risk for each location. Simulations integrate local climate profiles with experimentally derived biological parameters to assess outbreak timing and magnitude across geographically distinct settings (Chakraborty, Zigmond, et al., 2025; Couret & Benedict, 2014; Simoy et al., 2015; Yang et al., 2009).

### Spatiotemporal dynamics of egg drought tolerance in mosquito survival

Incorporating egg drought tolerance alongside thermal tolerance of other life stages revealed persistent mosquito populations and likely shifts in mosquito occurrence cycles over the coming decade (Figures 5, 6). We estimated early and late egg hatching probabilities for each location (six African locations of origin and six case study locations), offering refined insight into seasonal emergence patterns (Supplementary Figure 6, 8). In African contexts, rural regions exhibited a marked decline in egg hatching probability during the early months, with limited recovery projected for the post-monsoon period in the latter part of the year (Figure 5a). Urban populations showed consistently higher egg hatching probability compared to rural populations (Figure 5b), with late (delayed) egg hatching probability was notably higher in urban (0.89) than rural (0.64) habitats. Peak hatching, both in urban and rural areas, occurred during the wetter months, such as July to August (early) and September to October (late), while minimum hatching was observed during the drier months, spanning January to April (early) and February to May (late). Building on egg hatching probability and extending the analysis to other life stages survival rate, overall mosquito survival peaks followed egg hatching peaks (Figure 5), with mean survival probability higher in urban habitats (0.67) than in rural ones (0.5). Rural regions showed pronounced seasonal declines, especially from February to June (Figure 5d). In urban areas mosquito occurrence was typically consistent throughout the year in urban areas (Figure 5c); however, by autumn, occurrence levels in both urban and rural areas began to converge (Supplementary Figure 7). These urban-rural differences were most pronounced in early-century projections, highlighting the role of microclimatic buffering in sustaining vector persistence in urban habitats.

**Figure 5:**
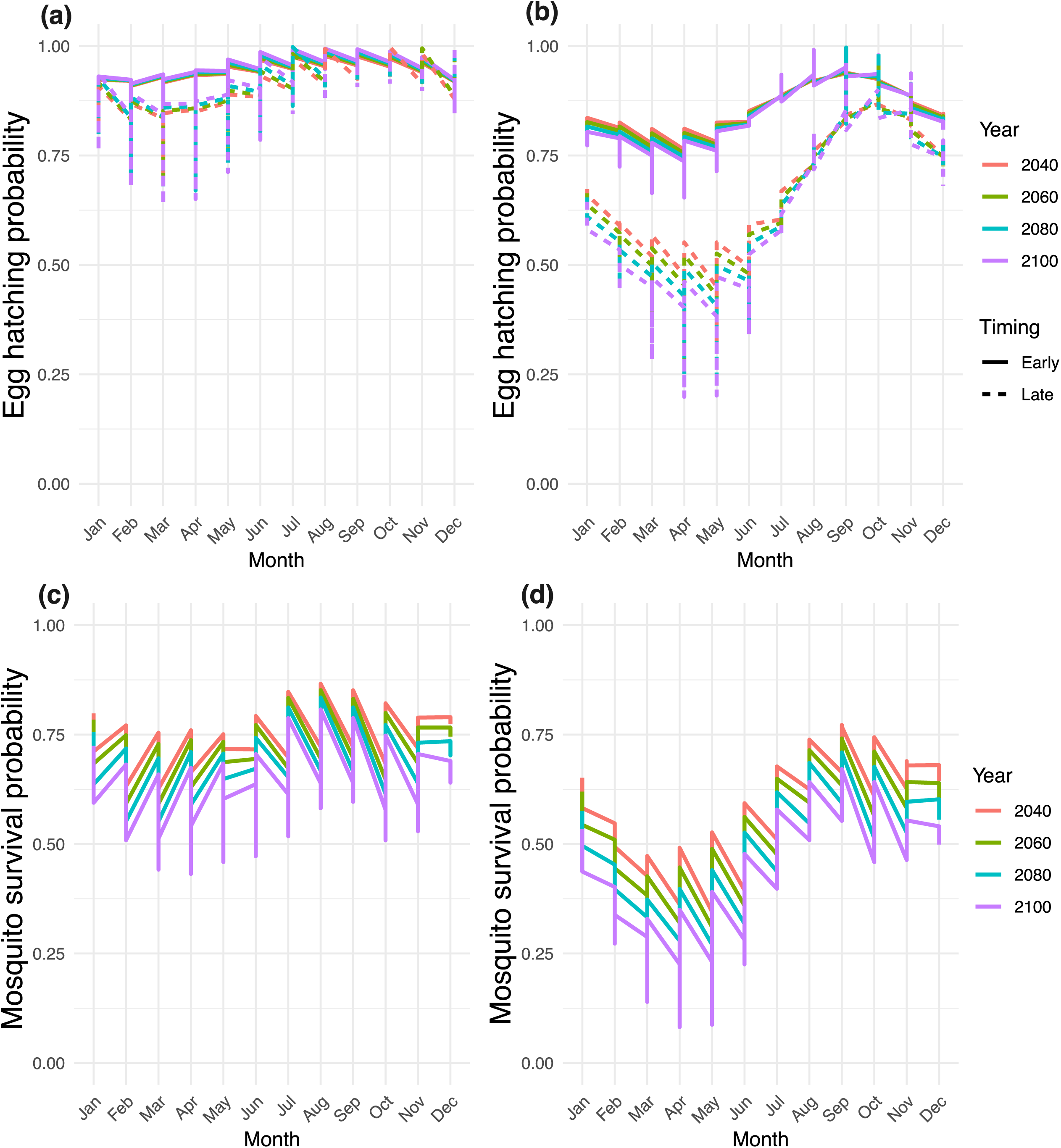
Projected egg hatching and survival for *Aedes aegypti* across urban and rural habitats under SSP3 future climate scenarios. Egg hatching probabilities for urban (a) and rural (b) populations and projected survival for first generation mosquitoes across the urban (c) and rural (d) habitats. Each colored line represents a different future decade, with line patterns corresponding to early (solid lines) or late (dotted lines) egg hatching probabilities. Model estimations are based on site-specific egg drought tolerance (a, b), which are combined with experimentally derived larval and adult thermal tolerance parameters to estimate the survival probabilities of the first generation (c, d). The jaggedness of the lines reflects inter-population variability, as the values represent averages across multiple urban (n = 3) and rural (n = 4) populations (see Supplementary Figures 6 and 7).

**Figure 6:**
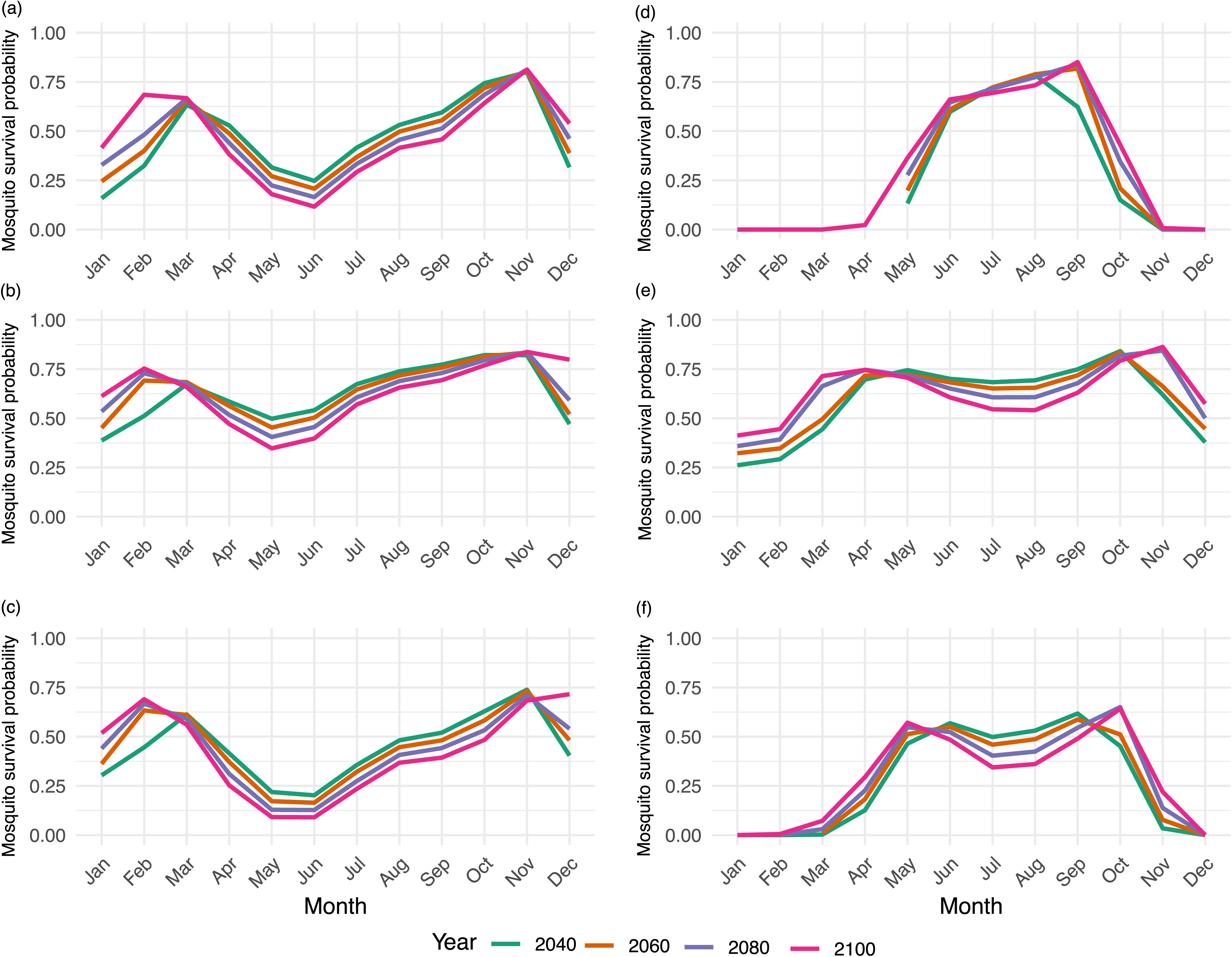
Projected monthly survival probabilities of mosquitoes across regions under SSP3 climate scenario. Simulated monthly survival probabilities of first-generation mosquitoes incorporating egg drought tolerance and larval/adult thermotolerance under SSP3 climate projections. Panels show projections for three Indian states, (a) Delhi, (b) Karnataka, and (c) Rajasthan, and three U.S. states, (d) Ohio, (e) Florida, and (f) Arizona. Colored lines represent different projection years: green (2040), orange (2060), blue (2080), and pink (2100).

Across all the locations in India and USA, egg hatching probabilities consistently revealed strong seasonality, typically peaking during the late summer and monsoon months (August to October) before declining in the winter (Supplementary Figure 8). While the three Indian locations and Florida maintained relatively high hatching probabilities year-round, more temperate or arid regions like Ohio and Arizona showed sharp reductions or near-zero probabilities during their respective cold or extremely dry seasons. Projected mosquito survival probabilities across all six case locations exhibited distinct geographic patterns and temporal shifts evident under future climate scenarios (Figure 6). In Indian locations, such as Delhi, Karnataka, and Rajasthan, survival probabilities exhibited a bimodal trend, peaking in March and again in October-November. Over the years, survival rates tended to increase in the early and late months (January-March, October-December), with modest decreases during the peak summer months (May-June). Rajasthan showed the most pronounced summer decline, with survival probabilities dropping below 0.2 threshold in June. In contrast, Ohio displayed highly seasonal survival, with negligible probabilities from January through March and from November onward. The summer months (June–September) showed a marked increase in survival probability, peaking at ∼0.9 in August. Additionally, slightly elevated survival probabilities in the early (April-May) and late (October-November) months by 2100 suggest a potential lengthening of the viable season. Florida exhibited relatively high year-round survival, with minor seasonal variation. Survival probabilities remained above 0.5 in nearly all months and increased slightly over time, particularly during the winter months (January-February and November-December), indicating a trend toward year-round suitability. In Arizona, a strong seasonal pattern was evident, with a peak from May to September. Notably, survival increased sharply in spring and fall months across time, and by 2100, survival exceeded 0.5 in both April and October. The summer months also saw slightly elevated probabilities in future years, reflecting an enhanced suitability. Overall, across all locations, later years (particularly 2100) were associated with increased survival during months that are currently less suitable, suggesting expanded temporal windows for mosquito persistence. However, some regions (notably Rajasthan and Delhi) may experience mid-year declines likely due to heat-related constraints, indicating potential nonlinear impacts of climatic warming.

## Discussion

Human expansion, rapid urbanization, and global warming have intensified interactions between humans and mosquitoes, driving the spread of mosquito-borne diseases and enabling mosquitoes to colonize and persist in regions once deemed inhospitable for their survival (de Souza & Weaver, 2024; Kache et al., 2022). This study highlights the critical role of *Aedes aegypti* egg desiccation resistance, examining how habitat-specific drought along with eco-climatic factors are driving mosquito population dynamics amid changing climatic conditions. Our findings confirm that *Ae. aegypti* egg viability declines with increasing saturation deficit (SD), especially after 12 weeks. Urban populations consistently exhibit higher egg drought tolerance than rural lineages under arid conditions, which were found to be associated with human specialization syndrome and bioclimatic normalities related to dry season temperature and wet season precipitation. We estimated *Ae. aegypti* survival probabilities under different SSP scenarios incorporating egg drought dynamics along with life stage-specific thermal tolerance, developmental timings, and reproductive output through a series of experimental and entomological transformations. These projections revealed egg hatching and population dynamics are significantly driven by climatic conditions and geographical locations. This study underscores that mosquito egg drought resilience is likely influenced by ancestral genomic background and is strongly shaped by human-associated habitats, which may intensify with increasing environmental fluctuations, raising significant concerns about future vector expansion in a warming world.

Temperature and humidity are crucial for egg viability (Brown et al., 2023; Lega et al., 2017); however, their combined effect, often quantified as SD, remains underexplored despite its direct influence on egg desiccation tolerance and, consequently, mosquito reproductive success and population dynamics (Faull & Williams, 2015; Holmes et al., 2022; Juliano et al., 2002). In Western Africa, SD rarely exceeds 20 mbar, while in subtropical and temperate zones, it can peak at 30 mbar during extreme heat. Our study, which spans a broader SD range (7 to 37 mbar), offers critical insights into how *Ae. aegypti* populations respond to prolonged desiccation stress across diverse environmental conditions. Egg viability declines sharply under prolonged high SD conditions, with a 75% reduction observed after three months at SD levels exceeding 25 mbar. However, more than 50% of eggs remain viable even after five months under lower SD conditions (below 25 mbar), highlighting a “safe SD zone” where desiccation stress is minimized. These trendstrends align with prior studies on other container-dwelling mosquito species, such as *Ae. notoscriptus,* which retained 9 - 13% viability after a year under lower SD levels (11 mbar) (Faull et al., 2016). Regional variations further highlight the role of local adaptation; for instance, eggs in northern Queensland retained over 88% viability under 14 mbar SD during the first two months but experienced a steep decline (2 - 15% hatching rates) after prolonged stress of one year (Faull & Williams, 2015). Overall, our findings support previous observations that eggs within humid areas distinctly allow *Ae. aegypti* to survive prolonged periods (Faull et al., 2016; Juliano et al., 2002; Sota & Mogi, 1992). Population-specific drought tolerance exhibited a clear distinction between urban and rural egg hatching at SD conditions exceeding 25 mbar. This relationship becomes particularly significant during prolonged drought conditions, which may explain the domesticity and anthropophilic tendencies of *Ae. aegypti* (de Souza & Weaver, 2024; Tchouassi et al., 2022). Urban environments offer abundant oviposition sites, such as water-holding artificial containers, stagnant pools, culverts, and sewers, providing mosquitoes with ample opportunities for egg deposition (Day, 2016; Renard et al., 2023). These artificial containers, often composed of anthropogenic pollutants, tend to be warmer due to substrate quality, and the urban heat island effect (Hsu et al., 2021; Tuholske et al., 2021) further exacerbates the impact of moist heat, which facilitates localized warming. In contrast, rural environments with natural oviposition sites, such as tree holes, may buffer against temperature extremes, benefiting from the shaded leafy canopies and the insulating effects of wood (Chakraborty, Zigmond, et al., 2025). Natural oviposition sites in rural areas, versus artificial containers and stagnant pools in urban settings likely shape the drought tolerance of mosquito eggs, as urban environments lack natural insulation and expose eggs to significantly higher thermal and desiccation stress.

Numerous studies have confirmed that human population density is a dominant ecological driver of *Ae. aegypti* distribution and spread (Abdalgader et al., 2022; Monaghan et al., 2018; Rose et al., 2023). Notably, varying human population densities across West African Sahel fundamentally shape mosquito-human associations, driving niche-specific mosquito adaptation that reflects their ancestral origin (Rose et al., 2020). The loss of natural breeding sites in tropical forested biomes, resulting from deforestation and climate change, is likely to drive mosquito populations towards human-altered environments with stable water sources, favorable microhabitats, and abundant opportunities for population growth (Laporta et al., 2023). Human population density and mosquito ancestry, linked to host preference, collectively explain the human specialization syndrome (Chakraborty, Zigmond, et al., 2025; Rose et al., 2020), a critical predictor of egg thermotolerance, which appeared significant for this study as well. Within this framework, ancestry emerges as the strongest determinant of egg drought resilience, followed by density, suggesting that specific loci linked to *‘Aaa’* ancestry and host-microhabitat dynamics may have a greater impact on genome-wide variation (Rose et al., 2020, 2023). Our results support this pattern of adaptation to urban setting, aligning with broader evidence across insect taxa, such as bed bugs, cattle ticks, tropical drosophilids, and phytophagous mites, showing genetic variations and local adaptations shaping reproductive responses to temperature fluctuations (Burrow, 2015; Moiroux et al., 2013; Mopper & Strauss, 2013; Wawrocka et al., 2015). However, substantial variation in egg drought tolerance can also reflect the impact of local rainfall ecology more than domestication or ancestry alone (Sota & Mogi, 1992; Valdez et al., 2018). *‘Aaf’* from prolonged dry regions of coastal Kenya survived longer egg desiccation with minimal mortality in laboratory setting, while the same sub-species from wetter inland regions of Kenya exhibited much shorter egg survival (Machado-Allison & Craig, 1972; Trpis, 1972). There is likewise a lot of variation within ‘*Aaa*’; colonies from coastal Kenya were highly sensitive to drying, whereas our result, from Senegal show high drought tolerance of ‘*Aaa*’ population (Mondet et al., 2005; Sylla et al., 2013). These observations caution against overgeneralizing the relationship between human association and drought tolerance, highlighting the influence of local eco-climatic context.

Complementing this genetic framework, bioclimatic factors, particularly the dynamics of temperature and precipitation-related variables during the dry and wet seasons, play a significant role in shaping female oviposition dynamics and subsequently regional egg drought tolerance (Chakraborty, Shah, et al., 2025; Venkataraman et al., 2023). Bioclimatic variables are key set of predictors often used to predict habitat suitability, disease risk, and long-term species persistence under current and future climate scenarios (Johnson et al., 2017; Lubinda et al., 2019; Njotto et al., 2024; Portilla Cabrera & Selvaraj, 2020). While previous work found minimal influence of local bioclimatic variables on egg thermotolerance (Chakraborty, Zigmond, et al., 2025), this study identifies a shift toward strong climate associations during the later stages of egg desiccation. Precipitation in the wettest month (bio13) and mean temperature of the driest quarter (bio9) emerged as key predictors of egg drought tolerance across timescales exceeding three months, reinforcing earlier findings that emphasize the combined influence of rainfall and heat accumulation (e.g., growing degree days) in supporting *Aedes spp.* proliferation (Iwamura et al., 2020; Johnson et al., 2017; Lubinda et al., 2019). Our models also found bio1 (annual mean temperature) to be influential, aligning with a previous study that identified bio1 as a major driver of future habitat suitability under climate change scenarios in Colombia (Portilla Cabrera & Selvaraj, 2020). While rainfall replenishes natural ephemeral pools and moisture necessary for egg and larval development (Liu & Yang, 2024; Singh et al., 2024), elevated dry-season temperatures accelerate evaporation, significantly reducing the persistence of suitable oviposition sites (Liu & Yang, 2024; Singh et al., 2024). These contrasting dynamics are particularly relevant given that ambient humidity levels are essential for egg viability, a pattern observed in ticks and mosquitoes (Mogi et al., 1996; Urbanski et al., 2010; Yoder et al., 2006). While our climate-based models revealed strong predictors of egg drought tolerance, the HSS framework, emphasizing the role of ancestry-mediated host preference and human population density, accounted for greater variation in the data. This finding underscores that even highly predictive climatic variables may contribute little to model performance when the primary ecological and evolutionary drivers are rooted in human–mosquito interactions.

Consistent with broader trends across insect taxa, recent studies highlight that population-specific variation in *Aedes* mosquito thermal performance is not merely regional, but is specifically shaped by local adaptation of life-history traits across diverse temperature gradients (Bong et al., 2021; Chakraborty, Zigmond, et al., 2025; Couper et al., 2024; Dennington et al., 2024; Rose et al., 2020). Such variations challenge the applicability of ‘one-size-fits-all’ approaches (Dennington et al., 2025), particularly as thermal optima and performance breadths diverge across populations. While previous results support the ‘hotter-is-better’ hypothesis, suggesting that populations adapted to higher temperatures exhibit greater fitness and broader performance curves (Alruiz et al., 2023; Dennington et al., 2025; Huey & Kingsolver, 1989), our simulation-based findings urge caution in generalizing this pattern across all ecological contexts when long-term viability of eggs with saturation deficit differences. Specifically, our models reveal that while elevated summer temperatures can accelerate mosquito development and reproduction, they may also reduce egg survival during overwintering and dry seasons, ultimately threatening long-term population viability, particularly in regions such as Ohio and Arizona. These patterns align more closely with the “jack-of-all-temperatures” hypothesis (also known as the specialist–generalist trade-off) (Huey & Kingsolver, 1989; Pawar et al., 2024), which posits that organisms cannot maximize performance across all conditions simultaneously. This is especially relevant when considering potential thermal and survival differences between *Ae. aegypti* forms like *‘Aaa’* and *‘Aaf’*. The extended egg and larval development times in urban settings may result from intensified intraspecific competition within abundant artificial breeding sites (Zahouli et al., 2016). Higher adult survival rates in urban environments may be attributable to consistent access to blood meals and reduced predation pressure (Goodman et al., 2018). Urban mosquitoes exhibited shorter oviposition intervals, likely due to increased host availability and higher biting opportunities in densely populated areas (Holmes et al., 2025). Although rural populations exhibited slightly higher egg production during peak reproductive months, the simulations projected that mosquito populations could re-establish within a year, with total population sizes substantially higher in urban areas, underscoring the heightened vector potential in urbanized environments. While studying the case sites alongside African locations, we noticed a “slow-burn” dynamic for mosquito population growth in Indian scenarios, where steady accumulation of viable eggs and favorable conditions gradually lead to a population surge that crosses an outbreak threshold. Noteworthy to mention, egg hatching probability peaks during the monsoon and late summer months (August-October) both in Indian and USA locations, identifying temperature and precipitation-triggered immersion as the primary drivers of *Ae. aegypti* population dynamics (Lega et al., 2017; Trpis, 1972). Delhi and Florida maintain year-round egg hatching as temperatures rarely drop below the critical 14 °C threshold, whereas Ohio and Arizona exhibit sharp seasonal declines to near-zero probability during winter or extreme dry seasons, marking the physiological limits of eggs for successful hatching (Eisen et al., 2014). Divergence in egg hatching across geographic locales suggests that anthropogenic climate warming may expand the “hatching windows” in temperate zones like Ohio, it could also push already arid regions like Rajasthan toward upper thermal thresholds that suppress egg viability (Sharma et al., 2025). Ultimately, our temperature-driven modeling framework captures shifts in fecundity, oviposition timing, and other seasonal bottlenecks that define *Ae. aegypti* persistence, where even small founder populations can trigger rapid expansion once environmental conditions become favorable.

Moving beyond thermal constraints, the integration of egg drought tolerance alongside temperature-dependent survival across life stages, our simulations revealed stark differences in mosquito population persistence. Under these multidimensional stressors, SSP3 (an emissions pathway marked by regional rivalry, low international cooperation, and limited mitigation efforts) emerges as the most plausible trajectory for future risk assessment (Riahi et al., 2017; Scafetta, 2024). The pronounced gap in hatching success and subsequent survival between urban and rural regions reflects the asymmetric impact of local climatic stressors, with urban areas offering more thermal and moisture refugia that allow the survival of mosquito populations. Early hatching (Week 1-8 after egg laying) may enable rapid population buildup under favorable conditions, while delayed hatching (Weeks 12-20 after egg laying) serves as a buffer during unfavorable periods (Faull & Williams, 2015). Moreover, historical data supported the validity of model projections, enhancing our capacity to anticipate future mosquito population surges and suggesting a persistent presence of mosquitoes, particularly in rapidly urbanizing regions (Njotto et al., 2024). The adaptive strategies in mosquito population surges are common across many regions globally and are believed to be primarily driven by temperature fluctuations (Lubinda et al., 2019; Portilla Cabrera & Selvaraj, 2020). However, our studies emphasize that humidity levels also have a significant impact. Egg drought tolerance refined our understanding of early and late hatching patterns, revealing greater resilience and adaptive potential of mosquito populations under urban environmental conditions. Furthermore, mosquito survival is closely aligned with periods of maximum egg hatching, which coincides with the wetter months, exacerbating their abundance (Newman et al., 2024). Our findings reinforce prior evidence that even modest climate shifts can have disproportionate impacts on vector ecology and further suggest that urban environmental conditions may increasingly alter mosquito occurrence trajectories (Johnson et al., 2017; Lippi et al., 2019; Wilke, Vasquez, et al., 2021). Although, we observed that egg survival and generational success between urban and rural areas appeared to converge toward the century’s end, potentially suggesting broader ecological transformations such as urban expansion, deforestation, and landscape alteration and anthropogenic warming (Norris, 2004; Ortiz et al., 2021). Despite these advances, some limitations remain, as our framework does not currently account for density-dependent feedbacks or track post-hatch development, which may lead to inaccurate estimates of cumulative mortality across life stages. Nevertheless, this predictive model serves as a valuable starting point, one that can be further refined by incorporating parameters, most importantly precipitation measures, from further studies.

Biome loss, rapid urbanization, pollution, and overexploitation are not only disrupting established ecosystems but also generating new human-associated niches that support the survival and reproduction of specific mosquito species. Our study highlights egg drought tolerance as a critical factor in understanding mosquito adaptability to changing environmental conditions. By examining egg-stage resilience under varying saturation deficits, we emphasize the extended drought tolerance of *Ae. aegypti* eggs and underscore how early-life traits influence population persistence across diverse habitats. These findings are particularly relevant as climate extremes become more frequent, suggesting the need for mosquito distribution and disease risk models to incorporate not only adult responses, but key developmental stages, such as the egg, that are critical to long-term survival under periods of low water availability. Understanding egg drought tolerance in the context of human specialization and climatic scenarios deepens our insight into the egg stage as a key to *Ae. aegypti* survival. This study highlights the importance of considering a range of factors, including desiccation stress, local climate, and the impact of human association, when predicting mosquito population dynamics and their range expansion.

## Acknowledgements

This study was partially supported by the National Institute of Allergy and Infectious Diseases of the National Institutes of Health under Award Numbers R01AI148551 and R21AI166633.

## Data availability

Zenodo. https://doi.org/10.5281/zenodo.18118396

## Supplementary Figures

**Supplementary Figure 1: Experimental setup for assessing egg viability across various moisture gradients over time**

A batch of 50–70 eggs were placed in 8 milliliter plastic vials (n = 10 vials/treatment), and the vials were housed inside desiccators maintained at one of three relative humidity (RH) levels: 33% (magnesium sulfate solution), 55% (calcium nitrate solution), or 75% (sodium chloride solution). Desiccators were placed inside climate-controlled incubators set at 23°C, 29°C, or 35°C (±1°C). Internal RH and temperature were verified using data loggers (ONSET HOBO). Eggs were stored under each of the nine temperature-humidity combinations and assessed for hatching after 1, 4, 8, 12, 16, and 20 weeks.

**Supplementary Figure 2: Temporal differences in egg hatching between urban and rural mosquito populations across Saturation Deficit levels**

Wilcoxon rank-sum tests assessing the effect of habitat (urban vs. rural) on egg hatching across varying time points and saturation deficit (SD) levels. Green cells indicate statistically significant differences between habitats, while white cells denote no significant difference.

**Supplementary Figure 3: Egg hatching threshold across urban and rural populations under varying saturation deficit conditions.**

Egg hatching thresholds across different populations over a 20-week desiccation period, with each panel representing a distinct population. The red-shaded panels indicate urban populations, while the blue-shaded panels represent rural populations. Each line corresponds to a specific saturation deficit (SD) level, derived from combinations of temperature (°C) and relative humidity (RH).

**Supplementary Figure 4: Egg hatching responses over time across a range of Saturation Deficit conditions**

Each panel represents a specific time point (in weeks) of egg exposure to varying saturation deficit (SD) levels. Points are colored by habitat type: rural (blue) and urban (red). This faceted scatter plot illustrates the general trend of declining egg hatching probability with increasing SD, highlighting how prolonged exposure to dry conditions reduces egg viability over time.

**Supplementary Figure 5: Time to 50% and 75% egg viability loss across saturation deficit levels**

Critical time points of egg viability loss are shown using bar plots, where brown bars indicate LT_50_ (week when 50% of eggs fail to hatch) and gray bars indicate LT_75_ (week when 75% fail to hatch), across three representative SD levels: 25.3, 26.83, and 37.66 mb. The aggregated egg hatching mean with 10-12 replicates with 50-75 eggs were used to generate the figure (Supplementary table ST3).

**Supplementary Figure 6: Projected egg hatching probabilities under future climate scenarios across six African locations.**

Line type denotes early (solid) and late (dashed) egg hatching periods and color indicates projection year for the locations: (a) Thies, (b) Kumasi, (c) Ngoye, (d) Kedougou, (e) Mindin, (f) Kintampo. Seasonal hatching profiles are derived from modeled relationships with saturation deficit (SD). Early hatching refers to eggs capable of surviving and hatching for up to eight weeks post-laying, while late hatching corresponds to survival extending to 16 weeks.

**Supplementary Figure 7: Projected mosquito survival probabilities across six African locations under SSP3 climate scenario.**

Line color denotes projection year of (a) Thies, (b) Kumasi, (c) Ngoye, (d) Kedougou, (e) Mindin, (f) Kintampo. Seasonal survival dynamics reflect downstream consequences of egg drought tolerance and temperature-based survival in subsequent life stages.

**Supplementary Figure 8: Egg hatching probabilities across selected study locations under future climate projections.**

Location-specific seasonal shifts in mosquito occurrence of (a) Delhi, (b) Karnataka, (c) Rajasthan, (d) Ohio, (e) Florida, and (f) Arizona. Line type represents early (solid) and late (dashed) egg hatching. Line color corresponds to projection year.

**Supplementary Table 1: Information on mosquito populations**

Collection localities and bioclimatic information about the *Aedes aegypt*i populations. Human population densities were calculated within a 20 km radius. Latitude and longitude coordinates represent the locations of egg ovitraps in the collection areas.

**Supplementary Table 2: Mean egg hatching responses of *Aedes aegypti* populations across saturation deficit (SD) and time.**

Average egg hatching rate and standard deviation for each mosquito population across different SD levels and time intervals.

**Supplementary Table 3: Egg hatching thresholds of *Aedes aegypti* populations under desiccation stress.**

The minimum time required to reduce hatching by 50% (LT_50_) and 75% (LT_75_) was estimated for each population using nested ANOVA models across multiple stress durations.

## References

Abdalgader, T., Pedersen, M., Ren, D., Sun, G., & Zhang, L. (2022). Trade-off between climatic and human population impacts on *Aedes aegypti* life history shapes its geographic distribution. Journal of Theoretical Biology, 535(110987), 110987.

Ajayi, O. M., Oyen, K. J., Davies, B., Finch, G., Piller, B. D., Harmeyer, A. A., Wendeln, K., Perretta, C., Rosendale, A. J., & Benoit, J. B. (2024). Egg hatching success is influenced by the time of thermal stress in four hard tick species. Journal of Medical Entomology, 61(1), 110–120.

Alruiz, J. M., Peralta-Maraver, I., Bozinovic, F., Santos, M., & Rezende, E. L. (2023). Temperature adaptation and its impact on the shape of performance curves in *Drosophila* populations. *Proceedings*. Biological Sciences, 290(1998), 20230507.

Arnau, J., Bono, R., Blanca, M. J., & Bendayan, R. (2012). Using the linear mixed model to analyze nonnormal data distributions in longitudinal designs. Behavior Research Methods, 44(4), 1224–1238.

Baty, F., Ritz, C., Charles, S., Brutsche, M., Flandrois, J.-P., & Delignette-Muller, M.-L. (2015). A toolbox for nonlinear regression in R: The package nlstools. Journal of Statistical Software, 66(5), 1–21.

Bong, L.-J., Tu, W.-C., & Neoh, K.-B. (2021). Interpopulation variations in life history traits and reproductive tactics in *Aedes aegypti*: A test on populations 50 km apart. Acta Tropica, 213(105750), 105750.

Bowler, K., & Terblanche, J. S. (2008). Insect thermal tolerance: what is the role of ontogeny, ageing and senescence? Biological Reviews of the Cambridge Philosophical Society, 83(3), 339–355.

Brown, J. J., Pascual, M., Wimberly, M. C., Johnson, L. R., & Murdock, C. C. (2023). Humidity - The overlooked variable in the thermal biology of mosquito-borne disease. Ecology Letters, 26(7), 1029–1049.

Burrow, H. M. (2015). Genetic aspects of cattle adaptation in the tropics. In The genetics of cattle (pp. 571–597). CABI.

Chakraborty, S., Shah, D., Dayal, D., Lutz, L., Wang, L., Garcia, J., Lefevre, G., Susanto, E., & Benoit, J. B. (2025). Parental exposure to wet and dry conditions shapes the viability and thermotolerance of eggs in *Aedes aegypti*. In bioRxivorg. 10.1101/2025.06.16.658957

Chakraborty, S., Zigmond, E., Shah, S., Dayal, D., Sylla, M., Akorli, J., Otoo, S., Rose, N. H., McBride, C. S., Armbruster, P. A., & Benoit, J. B. (2025). Thermal tolerance of *Aedes aegypti* mosquito eggs is associated with urban adaptation and human interactions. Journal of Thermal Biology, 131(104167), 104167.

Chu, E., Chakraborty, S., Benoit, J. B., & DeGennaro, M. (2022). Chapter 25: Water homeostasis and hygrosensation in mosquitoes. In Sensory ecology of disease vectors (pp. 655–682). Brill | Wageningen Academic.

Church, S. H., Donoughe, S., de Medeiros, B. A. S., & Extavour, C. G. (2019). Insect egg size and shape evolve with ecology but not developmental rate. Nature, 571(7763), 58–62.

Climate Change Knowledge Portal. (2025). https://climateknowledgeportal.worldbank.org/

Cnaan, A., Laird, N. M., & Slasor, P. (1997). Using the general linear mixed model to analyse unbalanced repeated measures and longitudinal data. Statistics in Medicine, 16(20), 2349–2380.

Couper, L. I., Farner, J. E., Lyberger, K. P., Lee, A. S., & Mordecai, E. A. (2024). Mosquito thermal tolerance is remarkably constrained across a large climatic range. In Proceedings of the Royal Society B: Biological Sciences. 10.1098/rspb.2023.2457

Couret, J., & Benedict, M. Q. (2014). A meta-analysis of the factors influencing development rate variation in *Aedes aegypti* (Diptera: Culicidae). BMC Ecology, 14(1), 3.

Day, J. F. (2016). Mosquito oviposition behavior and vector control. Insects, 7(4), 65.

Dennington, N. L., Grossman, M. K., Teeple, J. L., Johnson, L. R., Shocket, M. S., McGraw, E. A., & Thomas, M. B. (2025). Local adaptation of the mosquito vector, *Aedes aegypti*, and implications for predicting the effects of temperature and climate change on dengue transmission. In bioRxiv. 10.1101/2025.02.14.637444

Dennington, N. L., Grossman, M. K., Ware-Gilmore, F., Teeple, J. L., Johnson, L. R., Shocket, M. S., McGraw, E. A., & Thomas, M. B. (2024). Phenotypic adaptation to temperature in the mosquito vector, *Aedes aegypti*. Global Change Biology, 30(1), e17041.

de Souza, W. M., & Weaver, S. C. (2024). Effects of climate change and human activities on vector-borne diseases. Nature Reviews. Microbiology, 22(8), 476–491.

Eisen, L., Monaghan, A. J., Lozano-Fuentes, S., Steinhoff, D. F., Hayden, M. H., & Bieringer, P. E. (2014). The impact of temperature on the bionomics of *Aedes (Stegomyia) aegypti*, with special reference to the cool geographic range margins. Journal of Medical Entomology, 51(3), 496–516.

Estrada-Peña, A., Estrada-Sánchez, A., & de la Fuente, J. (2014). A global set of Fourier-transformed remotely sensed covariates for the description of abiotic niche in epidemiological studies of tick vector species. Parasites & Vectors, 7(1), 302.

Faull, K. J., Webb, C., & Williams, C. R. (2016). Desiccation survival time for eggs of a widespread and invasive Australian mosquito species, Aedes (Finlaya) notoscriptus (Skuse). Journal of Vector Ecology: Journal of the Society for Vector Ecology, 41(1), 55–62.

Faull, K. J., & Williams, C. R. (2015). Intraspecific variation in desiccation survival time of *Aedes aegypti* (L.) mosquito eggs of Australian origin. Journal of Vector Ecology: Journal of the Society for Vector Ecology, 40(2), 292–300.

Fick, S. E., & Hijmans, R. J. (2017). WorldClim 2: new 1-km spatial resolution climate surfaces for global land areas. International Journal of Climatology: A Journal of the Royal Meteorological Society, 37(12), 4302–4315.

Fischer, S., De Majo, M. S., Di Battista, C., & Campos, R. E. (2025). Effects of temperature and humidity on the survival and hatching response of diapausing and non-diapausing *Aedes aegypti* eggs. Journal of Insect Physiology, 161(104726), 104726.

Gibb, R., Colón-González, F. J., Lan, P. T., Huong, P. T., Nam, V. S., Duoc, V. T., Hung, D. T., Dong, N. T., Chien, V. C., Trang, L. T. T., Kien Quoc, D., Hoa, T. M., Tai, N. H., Hang, T. T., Tsarouchi, G., Ainscoe, E., Harpham, Q., Hofmann, B., Lumbroso, D., … Lowe, R. (2023). Interactions between climate change, urban infrastructure and mobility are driving dengue emergence in Vietnam. Nature Communications, 14(1), 8179.

Goodman, H., Egizi, A., Fonseca, D. M., Leisnham, P. T., & LaDeau, S. L. (2018). Primary blood-hosts of mosquitoes are influenced by social and ecological conditions in a complex urban landscape. Parasites & Vectors, 11(1), 218.

Guarneri, A. A., Lazzari, C., Diotaiuti, L., & Lorenzo, M. G. (2002). The effect of relative humidity on the behaviour and development of *Triatoma brasiliensis*. Physiological Entomology, 27(2), 142–147.

H. C., V., N., S. N., M. K., J., R. P., T., P., B., B., A., G., S., C., J., V. D., P., V., S., M., A., Shariff, M., S., R., & B. M., S. (2024). Unraveling Dengue Dynamics: In-Depth Epidemiological and Entomological Analyses in Bengaluru, India. Journal of Tropical Medicine, 2024, 1–7.

Holmes, C. J., & Benoit, J. B. (2019). Biological adaptations associated with dehydration in mosquitoes. Insects, 10(11), 375.

Holmes, C. J., Brown, E. S., Sharma, D., Nguyen, Q., Spangler, A. A., Pathak, A., Payton, B., Warden, M., Shah, A. J., Shaw, S., & Benoit, J. B. (2022). Bloodmeal regulation in mosquitoes curtails dehydration-induced mortality, altering vectorial capacity. Journal of Insect Physiology, 137(104363), 104363.

Holmes, C. J., Chakraborty, S., Ajayi, O. M., Uhran, M. R., Frigard, R., Stacey, C. L., Susanto, E. E., Chen, S.-C., Rasgon, J. L., DeGennaro, M., Xiao, Y., & Benoit, J. B. (2025). Multiple blood feeding bouts in mosquitoes allow for prolonged survival and are predicted to increase viral transmission during dry periods. iScience, 28(2), 111760.

Hsu, A., Sheriff, G., Chakraborty, T., & Manya, D. (2021). Disproportionate exposure to urban heat island intensity across major US cities. Nature Communications, 12(1), 2721.

Huey, R. B., & Kingsolver, J. G. (1989). Evolution of thermal sensitivity of ectotherm performance. Trends in Ecology & Evolution, 4(5), 131–135.

Iwamura, T., Guzman-Holst, A., & Murray, K. A. (2020). Accelerating invasion potential of disease vector *Aedes aegypti* under climate change. Nature Communications, 11(1), 2130.

Johnson, T. L., Haque, U., Monaghan, A. J., Eisen, L., Hahn, M. B., Hayden, M. H., Savage, H. M., McAllister, J., Mutebi, J.-P., & Eisen, R. J. (2017). Modeling the Environmental Suitability for *Aedes (Stegomyia) aegypti* and *Aedes (Stegomyia) albopictus* (Diptera: Culicidae) in the Contiguous United States. Journal of Medical Entomology, 54(6), 1605–1614.

Juliano, S. A., O’Meara, G. F., Morrill, J. R., & Cutwa, M. M. (2002). Desiccation and thermal tolerance of eggs and the coexistence of competing mosquitoes. Oecologia, 130(3), 458–469.

Juliano, S. A., & Philip Lounibos, L. (2005). Ecology of invasive mosquitoes: effects on resident species and on human health: Invasive mosquitoes. Ecology Letters, 8(5), 558–574.

Kache, P. A., Santos-Vega, M., Stewart-Ibarra, A. M., Cook, E. M., Seto, K. C., & Diuk-Wasser, M. A. (2022). Bridging landscape ecology and urban science to respond to the rising threat of mosquito-borne diseases. Nature Ecology & Evolution, 6(11), 1601–1616.

Kakarla, S. G., Bhimala, K. R., Kadiri, M. R., Kumaraswamy, S., & Mutheneni, S. R. (2020). Dengue situation in India: Suitability and transmission potential model for present and projected climate change scenarios. The Science of the Total Environment, 739(140336), 140336.

Kuznetsova, A., Brockhoff, P. B., & Christensen, R. H. B. (2017). LmerTest package: Tests in linear mixed effects models. Journal of Statistical Software, 82(13), 1–26.

Laporta, G. Z., Potter, A. M., Oliveira, J. F. A., Bourke, B. P., Pecor, D. B., & Linton, Y.-M. (2023). Global Distribution of *Aedes aegypti* and *Aedes albopictus* in a Climate Change Scenario of Regional Rivalry. Insects, 14(1), 49.

Lega, J., Brown, H. E., & Barrera, R. (2017). *Aedes aegypti* (Diptera: Culicidae) abundance model improved with relative humidity and precipitation-driven egg hatching. Journal of Medical Entomology, 54(5), 1375–1384.

Linde, T. V. D., Hewitt, P., Nel, A., & Westhuizen, M. (1990). The influence of different constant temperatures and saturation deficits on the survival of adult *Culex (Culex) theileri* Theobald (Diptera: Culicidae) in the laboratory. Journal of the Entomological Society of Southern Africa, 53, 57–63.

Lippi, C. A., Stewart-Ibarra, A. M., Loor, M. E. F. B., Zambrano, J. E. D., Lopez, N. A. E., Blackburn, J. K., & Ryan, S. J. (2019). Geographic shifts in *Aedes aegypti* habitat suitability in Ecuador using larval surveillance data and ecological niche modeling: Implications of climate change for public health vector control. PLoS Neglected Tropical Diseases, 13(4), e0007322.

Liu, Y., & Yang, X. (2024). Life cycle dynamics of mosquitoes under varied environmental conditions. Journal of Mosquito Research. 10.5376/jmr.2024.14.0015

Lubinda, J., Treviño C, J. A., Walsh, M. R., Moore, A. J., Hanafi-Bojd, A. A., Akgun, S., Zhao, B., Barro, A. S., Begum, M. M., Jamal, H., Angulo-Molina, A., & Haque, U. (2019). Environmental suitability for *Aedes aegypti* and *Aedes albopictus* and the spatial distribution of major arboviral infections in Mexico. Parasite Epidemiology and Control, 6(e00116), e00116.

Machado-Allison, C. E., & Craig, G. B., Jr. (1972). Geographic Variation in Resistance to Desiccation in *Aedes aegypti* and *A. atropalpus* (Diptera: Culicidae). Annals of the Entomological Society of America, 65(3), 542–547.

Maïga, H., Yamada, H., & Bimbile-Somda, N. S. (2017). Guidelines for routine colony maintenance of *Aedes* mosquito species. IAEA Physical and Chemical.

Martín, M. E., Estallo, E. L., Estrada, L. G., Matiz Enriquez, C., & Stein, M. (2025). Desiccation Tolerance of *Aedes aegypti* and *Aedes albopictus* Eggs of Northeastern Argentina Origin. Tropical Medicine and Infectious Disease, 10(4), 116.

McBride, C. S., Baier, F., Omondi, A. B., Spitzer, S. A., Lutomiah, J., Sang, R., Ignell, R., & Vosshall, L. B. (2014). Evolution of mosquito preference for humans linked to an odorant receptor. Nature, 515(7526), 222–227.

Meola, R. (1964). The influence of temperature and humidity on embryonic longevity in *Aedes aegypti*. Annals of the Entomological Society of America, 57(4), 468–472.

Mogi, M., Miyagi, I., Abadi, K., & Syafruddin. (1996). Inter- and intraspecific variation in resistance to desiccation by adult *Aedes (Stegomyia) spp*. (Diptera: Culicidae) from Indonesia. Journal of Medical Entomology, 33(1), 53–57.

Moiroux, J., Delava, E., Fleury, F., & van Baaren, J. (2013). Local adaptation of a *Drosophila* parasitoid: habitat-specific differences in thermal reaction norms. Journal of Evolutionary Biology, 26(5), 1108–1116.

Monaghan, A. J., Sampson, K. M., Steinhoff, D. F., Ernst, K. C., Ebi, K. L., Jones, B., & Hayden, M. H. (2018). The potential impacts of 21st century climatic and population changes on human exposure to the virus vector mosquito *Aedes aegypti*. Climatic Change, 146(3-4), 487–500.

Mondet, B., Diaïté, A., Ndione, J.-A., Fall, A. G., Chevalier, V., Lancelot, R., Ndiaye, M., & Ponçon, N. (2005). Rainfall patterns and population dynamics of *Aedes (Aedimorphus) vexans arabiensis*, Patton 1905 (Diptera: Culicidae), a potential vector of Rift Valley Fever virus in Senegal. Journal of Vector Ecology: Journal of the Society for Vector Ecology, 30(1), 102–106.

Mopper, S., & Strauss, S. Y. (Eds.). (2013). Genetic structure and local adaptation in natural insect populations: Effects of ecology, life history, and behavior. Springer. 10.1007/978-1-4757-0902-5

Newman, E. A., Feng, X., Onland, J. D., Walker, K. R., Young, S., Smith, K., Townsend, J., Damian, D., & Ernst, K. (2024). Defining the roles of local precipitation and anthropogenic water sources in driving the abundance of *Aedes aegypti*, an emerging disease vector in urban, arid landscapes. Scientific Reports, 14(1), 2058.

Njotto, L. L., Senyoni, W., Cronie, O., Alifrangis, M., & Stensgaard, A.-S. (2024). Quantitative modelling for dengue and *Aedes* mosquitoes in Africa: A systematic review of current approaches and future directions for Early Warning System development. PLoS Neglected Tropical Diseases, 18(11), e0012679.

Norris, D. E. (2004). Mosquito-borne diseases as a consequence of land use change. EcoHealth, 1(1), 19–24.

Ortiz, D. I., Piche-Ovares, M., Romero-Vega, L. M., Wagman, J., & Troyo, A. (2021). The impact of deforestation, urbanization, and changing land use patterns on the ecology of mosquito and tick-borne diseases in Central America. Insects, 13(1), 20.

Pawar, S., Huxley, P. J., Smallwood, T. R. C., Nesbit, M. L., Chan, A. H. H., Shocket, M. S., Johnson, L. R., Kontopoulos, D.-G., & Cator, L. J. (2024). Variation in temperature of peak trait performance constrains adaptation of arthropod populations to climatic warming. Nature Ecology & Evolution, 8(3), 500–510.

Portilla Cabrera, C. V., & Selvaraj, J. J. (2020). Geographic shifts in the bioclimatic suitability for *Aedes aegypti* under climate change scenarios in Colombia. Heliyon, 6(1), e03101.

Powell, J. R., Gloria-Soria, A., & Kotsakiozi, P. (2018). Recent History of *Aedes aegypti*: Vector Genomics and Epidemiology Records. Bioscience, 68(11), 854–860.

Renard, A., Pérez Lombardini, F., Pacheco Zapata, M., Porphyre, T., Bento, A., Suzán, G., Roiz, D., Roche, B., & Arnal, A. (2023). Interaction of human behavioral factors shapes the transmission of arboviruses by *Aedes* and *Culex* mosquitoes. Pathogens, 12(12), 1421.

Rezende, G. L., Martins, A. J., Gentile, C., Farnesi, L. C., Pelajo-Machado, M., Peixoto, A. A., & Valle, D. (2008). Embryonic desiccation resistance in *Aedes aegypti*: presumptive role of the chitinized serosal cuticle. BMC Developmental Biology, 8(1), 82.

Riahi, K., van Vuuren, D. P., Kriegler, E., Edmonds, J., O’Neill, B. C., Fujimori, S., Bauer, N., Calvin, K., Dellink, R., Fricko, O., Lutz, W., Popp, A., Cuaresma, J. C., Kc, S., Leimbach, M., Jiang, L., Kram, T., Rao, S., Emmerling, J., … Tavoni, M. (2017). The Shared Socioeconomic Pathways and their energy, land use, and greenhouse gas emissions implications: An overview. Global Environmental Change: Human and Policy Dimensions, 42, 153–168.

Rose, N. H., Badolo, A., Sylla, M., Akorli, J., Otoo, S., Gloria-Soria, A., Powell, J. R., White, B. J., Crawford, J. E., & McBride, C. S. (2023). Dating the origin and spread of specialization on human hosts in *Aedes aegypti* mosquitoes. eLife, 12, e83524.

Rose, N. H., Sylla, M., Badolo, A., Lutomiah, J., Ayala, D., Aribodor, O. B., Ibe, N., Akorli, J., Otoo, S., Mutebi, J.-P., Kriete, A. L., Ewing, E. G., Sang, R., Gloria-Soria, A., Powell, J. R., Baker, R. E., White, B. J., Crawford, J. E., & McBride, C. S. (2020). Climate and urbanization drive mosquito preference for humans. Current Biology: CB, 30(18), 3570–3579.e6.

Scafetta, N. (2024). Impacts and risks of “realistic” global warming projections for the 21st century. Geoscience Frontiers, 15(2), 101774.

Sharma, G., Khan, Z., Das, D., Singh, S., Singh, S., Kumar, M., Tiwari, R. R., & Sarma, D. K. (2025). Thermal influence on development and morphological traits of *Aedes aegypti* in central India and its relevance to climate change. Parasites & Vectors, 18(1), 279.

Simoy, M. I., Simoy, M. V., & Canziani, G. A. (2015). The effect of temperature on the population dynamics of *Aedes aegypti*. Ecological Modelling, 314, 100–110.

Singh, H., Akhtar, N., & Gupta, S. K. (2024). Biology of mosquitoes. In Mosquitoes (pp. 141–163). Springer Nature Singapore.

Sota, T., & Mogi, M. (1992). Interspecific variation in desiccation survival time of *Aedes (Stegomyia)* mosquito eggs is correlated with habitat and egg size. Oecologia, 90(3), 353–358.

Sukhralia, S., Verma, M., Gopirajan, S., Dhanaraj, P. S., Lal, R., Mehla, N., & Kant, C. R. (2019). From dengue to Zika: the wide spread of mosquito-borne arboviruses. European Journal of Clinical Microbiology & Infectious Diseases: Official Publication of the European Society of Clinical Microbiology, 38(1), 3–14.

Sylla, M., Ndiaye, M., & Black, W. C. (2013). *Aedes* species in treeholes and fruit husks between dry and wet seasons in southeastern Senegal. Journal of Vector Ecology: Journal of the Society for Vector Ecology, 38(2), 237–244.

Tchouassi, D. P., Agha, S. B., Villinger, J., Sang, R., & Torto, B. (2022). The distinctive bionomics of *Aedes aegypti* populations in Africa. Current Opinion in Insect Science, 54(100986), 100986.

Trpis, M. (1972). Dry season survival of *Aedes aegypti* eggs in various breeding sites in the Dar es Salaam area, Tanzania. Bulletin of the World Health Organization, 47(3), 433–437.

Tuholske, C., Caylor, K., Funk, C., Verdin, A., Sweeney, S., Grace, K., Peterson, P., & Evans, T. (2021). Global urban population exposure to extreme heat. Proceedings of the National Academy of Sciences of the United States of America, 118(41), e2024792118.

Urbanski, J. M., Benoit, J. B., Michaud, M. R., Denlinger, D. L., & Armbruster, P. (2010). The molecular physiology of increased egg desiccation resistance during diapause in the invasive mosquito, Aedes albopictus. Proceedings. Biological Sciences, 277(1694), 2683–2692.

Valdez, L. D., Sibona, G. J., & Condat, C. A. (2018). Impact of rainfall on *Aedes aegypti* populations. Ecological Modelling, 385, 96–105.

Varamballi, P., Babu N, N., Mudgal, P. P., Shetty, U., Jayaram, A., Karunakaran, K., Arumugam, S., & Mukhopadhyay, C. (2024). Spatial heterogeneity in the potential distribution of *Aedes* mosquitoes in India under current and future climatic scenarios. Acta Tropica, 260(107403), 107403.

Venkataraman, K., Shai, N., Lakhiani, P., Zylka, S., Zhao, J., Herre, M., Zeng, J., Neal, L. A., Molina, H., Zhao, L., & Vosshall, L. B. (2023). Two novel, tightly linked, and rapidly evolving genes underlie *Aedes aegypti* mosquito reproductive resilience during drought. eLife, 12(e80489). 10.7554/eLife.80489

Wawrocka, K., Balvín, O., & Bartonička, T. (2015). Reproduction barrier between two lineages of bed bug (*Cimex lectularius*) (Heteroptera: Cimicidae). Parasitology Research, 114(8), 3019–3025.

Weaver, S. C., Charlier, C., Vasilakis, N., & Lecuit, M. (2018). Zika, Chikungunya, and other emerging vector-borne viral diseases. Annual Review of Medicine, 69(1), 395–408.

Wilke, A. B. B., Benelli, G., & Beier, J. C. (2021). Anthropogenic changes and associated impacts on vector-borne diseases. Trends in Parasitology, 37(12), 1027–1030.

Wilke, A. B. B., Vasquez, C., Carvajal, A., Moreno, M., Fuller, D. O., Cardenas, G., Petrie, W. D., & Beier, J. C. (2021). Urbanization favors the proliferation of *Aedes aegypti* and *Culex quinquefasciatus* in urban areas of Miami-Dade County, Florida. Scientific Reports, 11(1), 22989.

Yadav, P. D., Malhotra, B., Sapkal, G., Nyayanit, D. A., Deshpande, G., Gupta, N., Padinjaremattathil, U. T., Sharma, H., Sahay, R. R., Sharma, P., & Mourya, D. T. (2019). Zika virus outbreak in Rajasthan, India in 2018 was caused by a virus endemic to Asia. Infection, Genetics and Evolution: Journal of Molecular Epidemiology and Evolutionary Genetics in Infectious Diseases, 69, 199–202.

Yang, H. M., Macoris, M. L. G., Galvani, K. C., Andrighetti, M. T. M., & Wanderley, D. M. V. (2009). Assessing the effects of temperature on the population of *Aedes aegypti*, the vector of dengue. Epidemiology and Infection, 137(8), 1188–1202.

Yoder, J. A., Benoit, J. B., Rellinger, E. J., & Tank, J. L. (2006). Developmental profiles in tick water balance with a focus on the new Rocky Mountain spotted fever vector, *Rhipicephalus sanguineus*. Medical and Veterinary Entomology, 20(4), 365–372.

Zahouli, J. B. Z., Utzinger, J., Adja, M. A., Müller, P., Malone, D., Tano, Y., & Koudou, B. G. (2016). Oviposition ecology and species composition of *Aedes spp.* and *Aedes aegypti* dynamics in variously urbanized settings in arbovirus foci in southeastern Côte d’Ivoire. Parasites & Vectors, 9(1), 523.

