## Supplemental methods for "Drought tolerance of *Aedes aegypti* mosquito eggs is influenced by adaptation to local climate conditions and associations with humans"

**Supplementary methods:**

**Part 1. Modeling temperature-dependent mosquito life cycle and population dynamics**

The local monthly average temperature was collected from an open-access website ("Climate Change Knowledge Portal," 2025). Due to the influence of varying environmental pressures across mosquito developmental stages, we used a stage-structured model to represent mosquito population dynamics under fluctuating environmental conditions. The model includes five distinct life stages: E (eggs), L (larvae and pupae), $A_1$ (newly emerged adults in their first gonotrophic cycle before oviposition), $A_2$ (female adults in their subsequent cycles before oviposition), and $A_3$ (ovipositing adults in subsequent cycles). The model incorporates biologically realistic time delays and temperature-dependent mortality, reflecting how environmental fluctuations impact developmental transitions. The delay differential equations of the stage-structured model track stage-specific changes over time, with key transitions governed by developmental delays ($\tau_{1(t)}$ and $\tau_{2(t)}$) and stage-specific death rates for eggs ($d_e$), larvae/pupae ($d_l$), and adults ($d_a$). The average number of eggs laid per female per day (e(t)) and successful egg hatch rates ($B_1$(A_3, t), $B_2$(A_2, t)) are also temperature dependent. Adult mosquitoes progress through three physiological stages post-emergence: $A_1$ represents females preparing for their first oviposition after eclosion, $A_2$ accounts for resting individuals between cycles, and $A_3$ includes mosquitoes post first oviposition cycle. Each adult stage contributes to egg production, with transitions governed by oviposition rate ($\alpha_1$ and $\alpha_2$), and $\alpha_3$ is the average duration of oviposition in subsequent cycles. Environmental stress, especially elevated saturation deficits, affects mortality and may delay transitions, reflecting the ecological reality that dehydration or thermal extremes influence reproductive timing and survival.

$$E' = B_1(A_3(t), t) + B_2(\alpha_1 A_1(t), t) - d_e(t)E(t) - e^{\int_{t-\tau_1(t)}^{t} d_e(s)ds} E\big(t - \tau_1(t)\big);$$

$$L' = e^{\int_{t-\tau_1(t)}^{t} d_e(s)ds} E(t - \tau_1(t)) - d_l L(t) - e^{\int_{t-\tau_2(t)}^{t} d_l(s)ds} L(t - \tau_2(t));$$

$$A_1' = e^{\int_{t-\tau_2(t)}^{t} d_l(s)ds} L(t - \tau_2(t)) - (d_a(t) + \alpha_1)A_1(t);$$

$$A_2' = \alpha_1 A_1(t) + \alpha_3 A_3(t) - (d_a + \alpha_2(t))A_2(t);$$

$$A_3' = \alpha_2(t)A_2(t) - (d_a + \alpha_3)A_3(t).$$

| Parameter | Description | Range / Notes | References |
|---|---|---|---|
| $B_1(\cdot,t)$, $B_2(\cdot,t)$ | Average number of eggs successfully hatched | - | (Chakraborty et al., 2024) |
| e(t) | Average number of eggs laid per female mosquito | 50-200 | (Chakraborty et al., 2024; Holmes et al., 2025) |
| $d_e(t)$ | Death rate of eggs (environmental effects) | 0-1 | (Chakraborty et al., 2024) |
| $d_l(t)$ | Death rate for larvae and pupae | 0-1 | (Chakraborty et al., 2024) |
| $d_a(t)$ | Death rate for adults | 0-1 | (Chakraborty et al., 2024) |
| $1/\alpha_1$ | Duration of the first oviposition cycle (post-eclosion) | 14 days | (Yang et al., 2009) |
| $1/\alpha_2$ | Duration of other (successive) oviposition cycles | 3-40 days | (Yang et al., 2009) |
| $\alpha_3$ | Duration of subsequent oviposition | 3-14 days | |
| $\tau_{1(t)}$ | Developmental delay from egg to larva/pupa | 0-55 days | (Couret and Benedict, 2014; Simoy et al., 2015) |
| $\tau_{2(t)}$ | Developmental delay from larva/pupa to adult | 0-60 days | (Couret and Benedict, 2014; Simoy et al., 2015) |

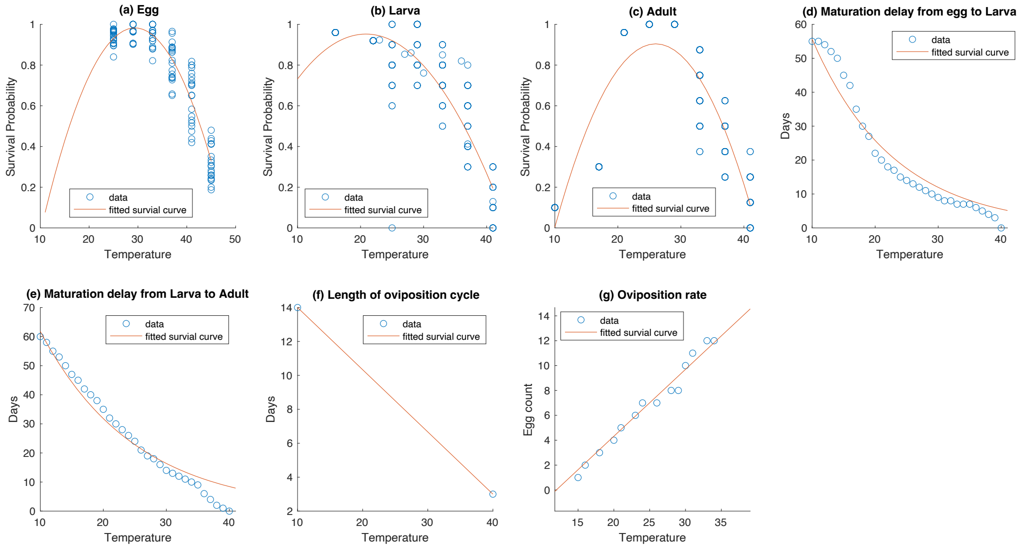


Life stage-specific survival probability, developmental delays, and gonotrophic cycles, including egg production rate, were modeled as a piecewise function of temperature (T, in °C):

**(a) Egg survival rate, (1-$d_{e(t)}$):** (Chakraborty et al., 2024)

- 25°C ≤ T ≤ 45°C ⟶ Survival follows a quadratic curve = -0.0028x2 + 0.1647x - 1.383

**(b) Larval survival rate, (1-$d_{l(t)}$):** (Chakraborty et al., 2024)

- T < 10°C ⟶ Survival = 0 (no survival)
- 10°C ≤ T < 16°C ⟶ Survival increases linearly = 0.15 × T - 1.5
- 16°C ≤ T ≤ 41°C ⟶ Survival follows a quadratic curve = -0.0019 × $T^2$ + 0.0775 × T + 0.1459
- T > 41°C ⟶ Survival = 0

**(c) Adult survival rate (1/$d_{a(t)}$):** (Chakraborty et al., 2024)

- T < 10°C ⟶ Survival = 0
- 10°C ≤ T < 21°C ⟶ Survival increases linearly = 0.0723 × T - 0.7045

- 21°C ≤ T ≤ 41°C → Survival follows a quadratic curve = -0.0029 × T² + 0.1375 × T - 0.6153
- T > 41°C → Survival = 0

**(d and e) Development delays function ($\tau_{1(t)}$ and $\tau_{2(t)}$):** (Chakraborty et al., 2024; Couret and Benedict, 2014; Simoy et al., 2015)

Development time (from egg to larvae and larvae to adult) was modeled using an exponential decay function, using $a \cdot e^{-b(T-10)}$, for T>10 °C, where T is temperature (°C), a and b are estimated parameters from nonlinear least squares fitting. Development was assumed to cease at temperatures ≤10°C (development time = ∞). Using experimental datasets and datasets extracted from the published studies, we fitted this exponential model and generated predicted developmental durations. These predictions were later used to infer life-cycle timing under varied climate scenarios.

(f) **Length of oviposition cycle:** (Yang et al., 2009) The duration of the first oviposition cycle, i.e., from adult emergence (eclosion) to completion of the egg-laying event, was assumed to be 14 days at 10°C, with progressively shorter durations at higher temperatures, reaching as few as 3 days at 41°C. The linear model =14+(3-14) × (x-10) / (41-10)

(g) **Daily egg production:** (Yang et al., 2009)

- T < 10°C → Egg production = 0
- 10°C ≤ T < 40°C → Egg production follows a quadratic curve = -0.000066 × T² + 0.579 × T - 7.387.
- T > 40°C → Egg production = 0

**Average number of eggs successfully hatched:** duration of gonotrophic cycles × daily egg production number, i.e., $B_1(A_3(t), t)$ = duration of the first gonotrophic cycles × daily egg production number, and $B_2$ $(\alpha_1 A_1 t(t)$ = duration of the successive gonotrophic cycles × daily egg production number

**Part 2. Inclusion of Relative Humidity (RH) to predict future mosquito survival probability**

Historical and projected future data for temperature and relative humidity were collected from the website ("Climate Change Knowledge Portal," 2025). There was inconsistency in the relative humidity data, so using historical data for RH and temperature, we predicted a temperature and RH for each month to cross-validate the website-generated future RH data. We predicted future RH for each SSP condition, using - $RH_h/T_h = (RH_f - RH_p)/(T_f - T_p)$ , where h = historic data; f = future data; p = model-predicted data. Fourier series decomposition (or a truncated Fourier series) within a nonlinear least square ('nls') regression model was used to capture the seasonal variation in temperature and relative humidity across months.

**2.1. Temperature and RH were converted to Saturation Deficit:** (Holmes and Benoit, 2019; Linde et al., 1990)

$$\left(\left(610.7 * \left(10^{\left(\frac{7.5*T}{237.2+T}\right)}\right)\right) * \frac{\frac{100-RH}{100}}{100}\right)$$

### 2.2. Calculation of early egg hatch and late egg hatch:

To reflect the observed long-term viability of *Aedes aegypti* eggs in our modeling framework, we calculated early and late egg hatching by considering hatching from the preceding two and four months, respectively. Experimental results demonstrated that *Aedes aegypti* eggs can survive for up to five months. To incorporate this biological constraint into our climate-based modeling framework, we calculated two metrics, i.e., early egg hatching (based on the current and previous month) and late egg hatching (based on the current and previous three months). The predicted probability of egg hatching in month $i$ was calculated as a weighted average of regression-based estimates from the relevant months, weighted by the number of days in each month ($d_i$):

Early egg hatching model:

$$y_i = \left(\frac{d_i}{d_i+d_{i-1}}\right)(m_1 x_i + c_1) + \left(\frac{d_{i-1}}{d_i+d_{i-1}}\right)(m_1 x_{i-1} + c_1)$$

Late egg hatching model:

$$y_i = \sum_{j=0}^{3}\left(\frac{d_{i-j}}{\sum_{k=3}^{3} d_{i-k}}\right)(m_2 x_{i-j} + c_2)$$

Where,

$y_i$: Predicted probability of egg hatching in month i;

$x_i$: Saturation deficit (SD)for month i ;

$m_1, c_1$: Regression slope and intercept from early egg hatching experimental data;

$m_2, c_2$: Slope and intercept from late egg hatching data;

$d_i$: Number of days in month i ;

**2.3. Entomological transformations of egg hatching-** To account for the nonlinear influence of temperature on egg survival, we applied a biologically derived transformation that scales the early and late hatching probabilities based on average monthly temperature. Each temperature value was mapped to a corresponding scaling coefficient representing relative egg survival potential. Final early and late hatching estimates were obtained by multiplying the predicted values by these scale factors. The overall monthly egg survival was calculated as the average of the scaled early and late hatching probabilities.

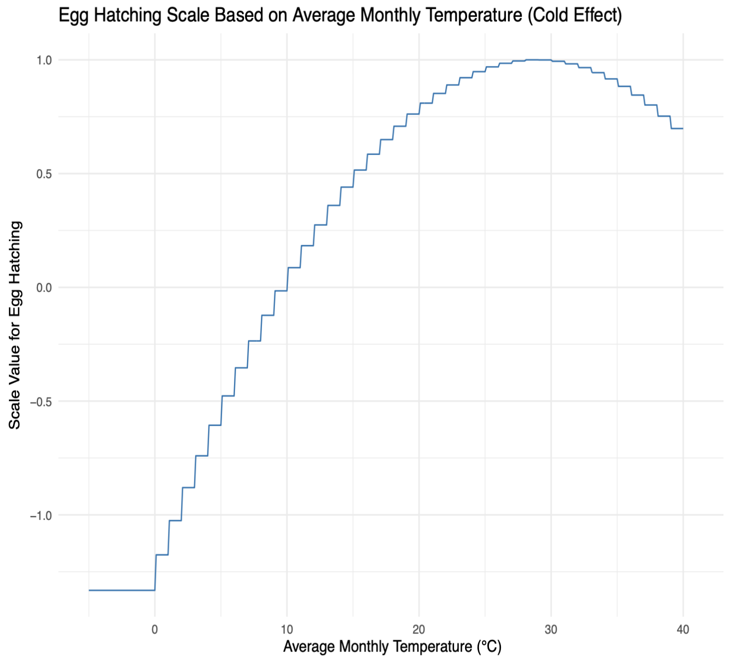


**2.4. Modeling mosquito survival probability:** To estimate the overall survival probability of the first mosquito generation, we multiplied egg drought tolerance with survival probabilities derived from larval and adult thermal tolerance. This integrative approach captured how stage-specific environmental limits jointly determine the likelihood of successful population establishment at a given location.

**References**


Chakraborty, S., Zigmond, E., Shah, S., Sylla, M., Akorli, J., Otoo, S., Rose, N.H., McBride, C.S., Armbruster, P.A., Benoit, J.B., 2024. Thermal tolerance of mosquito eggs is associated with urban adaptation and human interactions. bioRxivorg. https://doi.org/10.1101/2024.03.22.586322

Climate Change Knowledge Portal [World Bank], 2025. URL https://climateknowledgeportal.worldbank.org/ (accessed 6.2.25).

Couret, J., Benedict, M.Q., 2014. A meta-analysis of the factors influencing development rate variation in *Aedes aegypti* (Diptera: Culicidae). BMC Ecol. 14, 3.

Holmes, C.J., Benoit, J.B., 2019. Biological adaptations associated with dehydration in mosquitoes. Insects 10, 375.

Holmes, C.J., Chakraborty, S., Ajayi, O.M., Uhran, M.R., Frigard, R., Stacey, C.L., Susanto, E.E., Chen, S.-C., Rasgon, J.L., DeGennaro, M., Xiao, Y., Benoit, J.B., 2025. Multiple blood feeding bouts in mosquitoes allow for prolonged survival and are predicted to increase viral transmission during dry periods. iScience 28, 111760.

Linde, T.V.D., Hewitt, P., Nel, A., Westhuizen, M., 1990. The influence of different constant temperatures and saturation deficits on the survival of adult *Culex (Culex) theileri* Theobald (Diptera: Culicidae) in the laboratory. Journal of the Entomological Society of Southern Africa 53, 57–63.

Simoy, M.I., Simoy, M.V., Canziani, G.A., 2015. The effect of temperature on the population dynamics of *Aedes aegypti*. Ecol. Modell. 314, 100–110.

Yang, H.M., Macoris, M.L.G., Galvani, K.C., Andrighetti, M.T.M., Wanderley, D.M.V., 2009. Assessing the effects of temperature on the population of *Aedes aegypti*, the vector of dengue. Epidemiol. Infect. 137, 1188–1202.
