## Supplemental Figure for "Drought tolerance of *Aedes aegypti* mosquito eggs is influenced by adaptation to local climate conditions and associations with humans"

Supplementary Figure 1: Experimental setup for assessing egg viability across various moisture gradients over time.

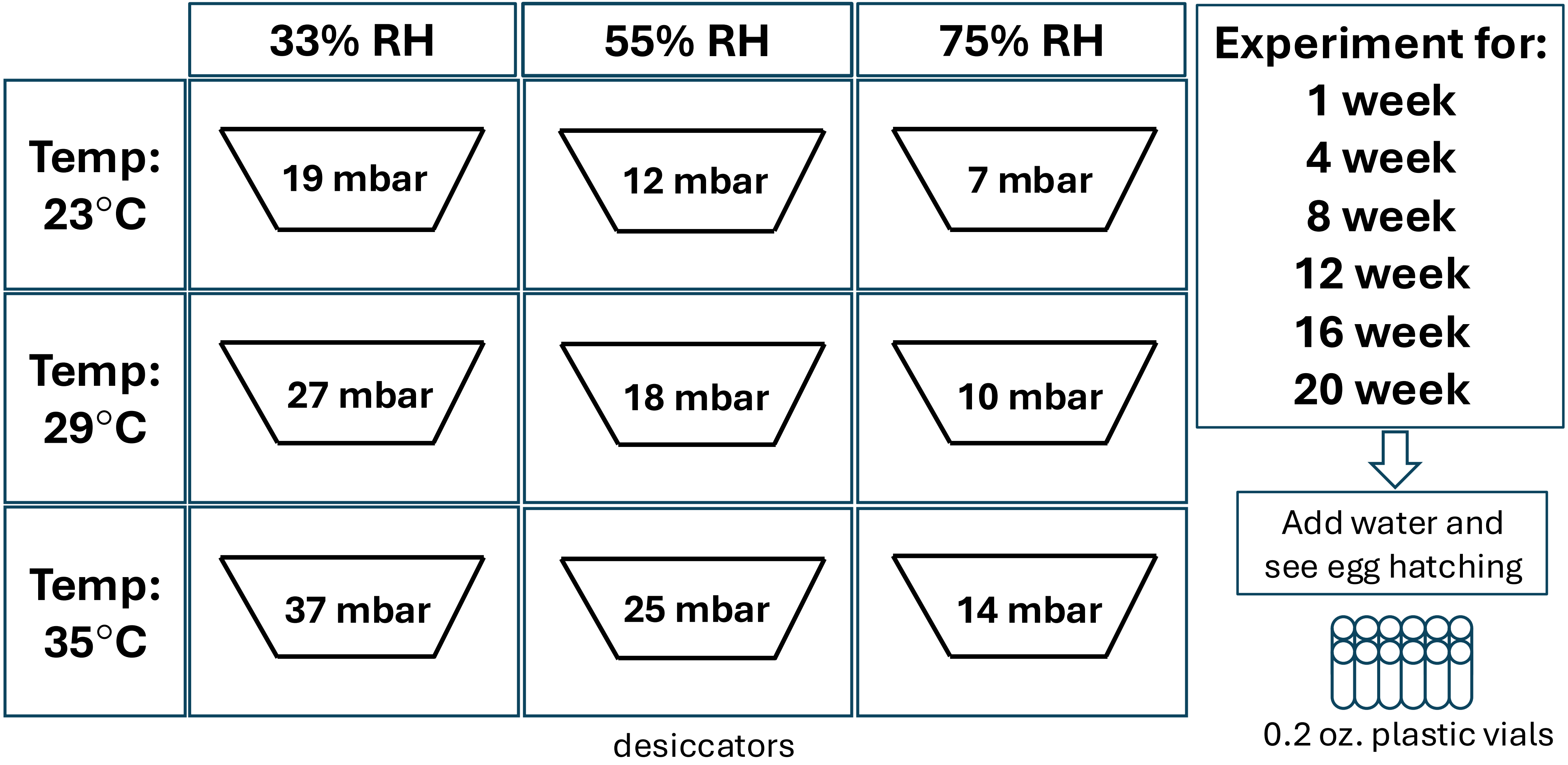

Supplementary Figure 2: Temporal differences in egg hatching between urban and rural mosquito populations across Saturation Deficit levels.

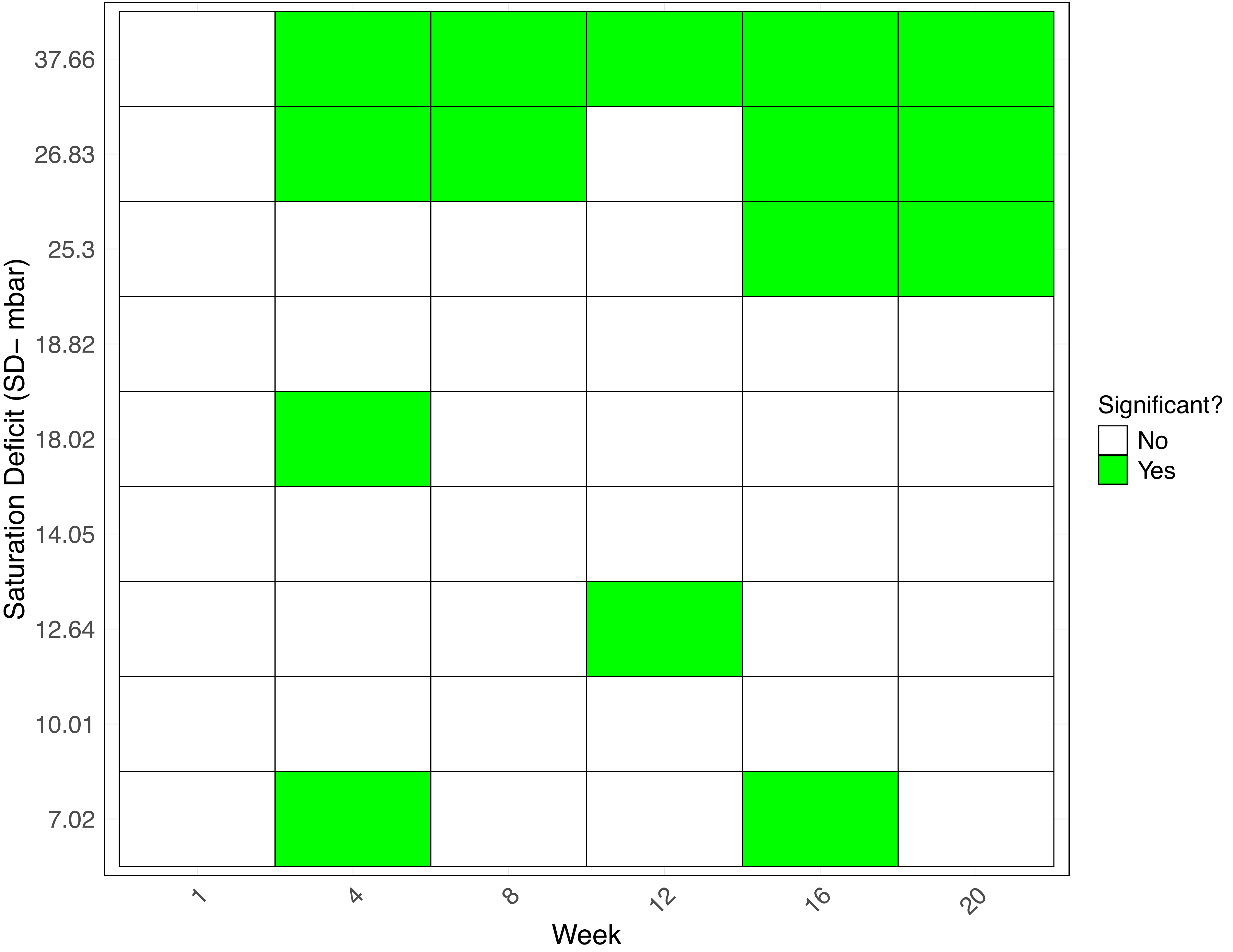

Supplementary Figure 3: Egg hatching threshold across urban and rural populations under varying saturation deficit conditions.

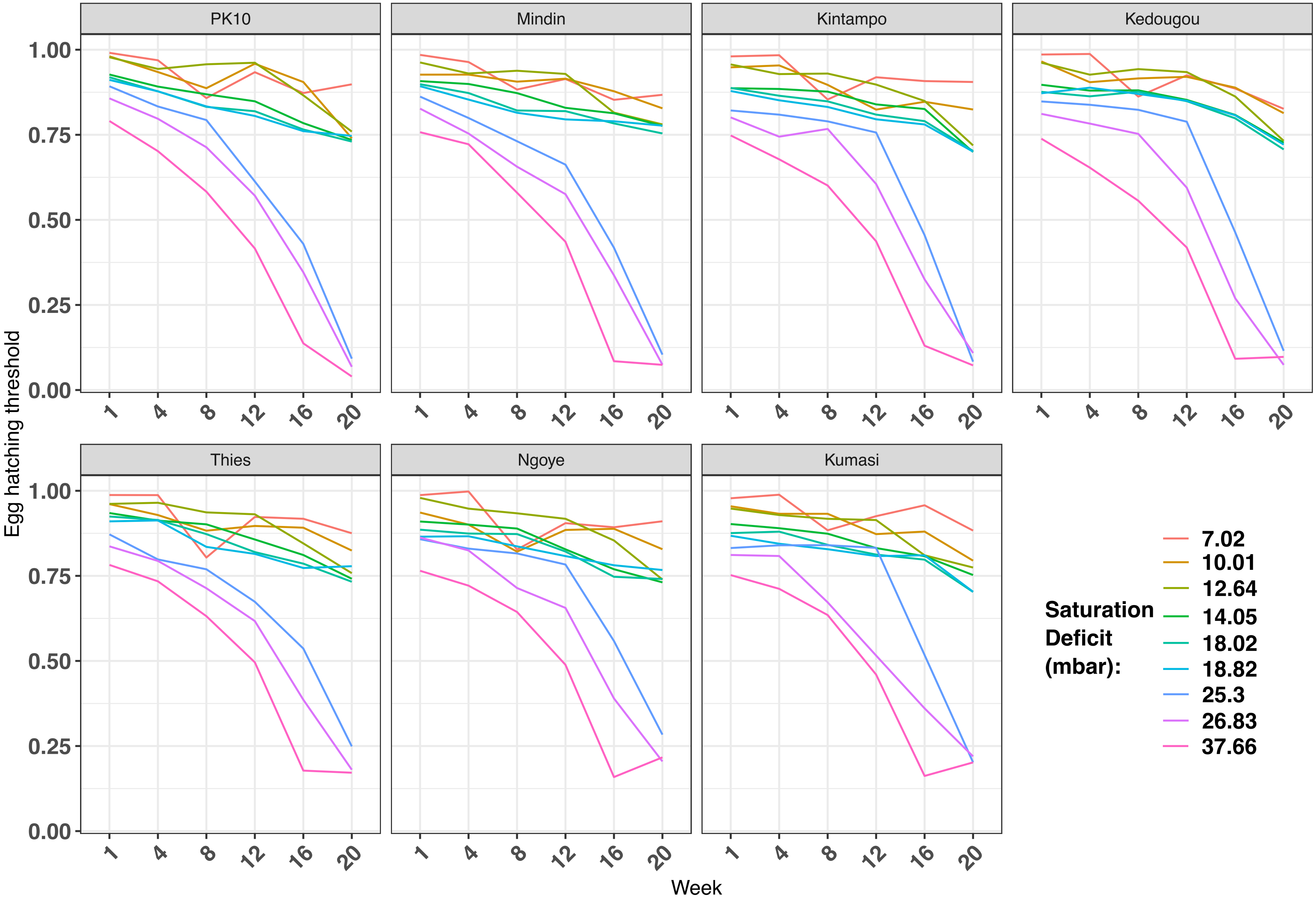

Supplementary Figure 4: Egg hatching responses over time across a range of Saturation Deficit conditions.

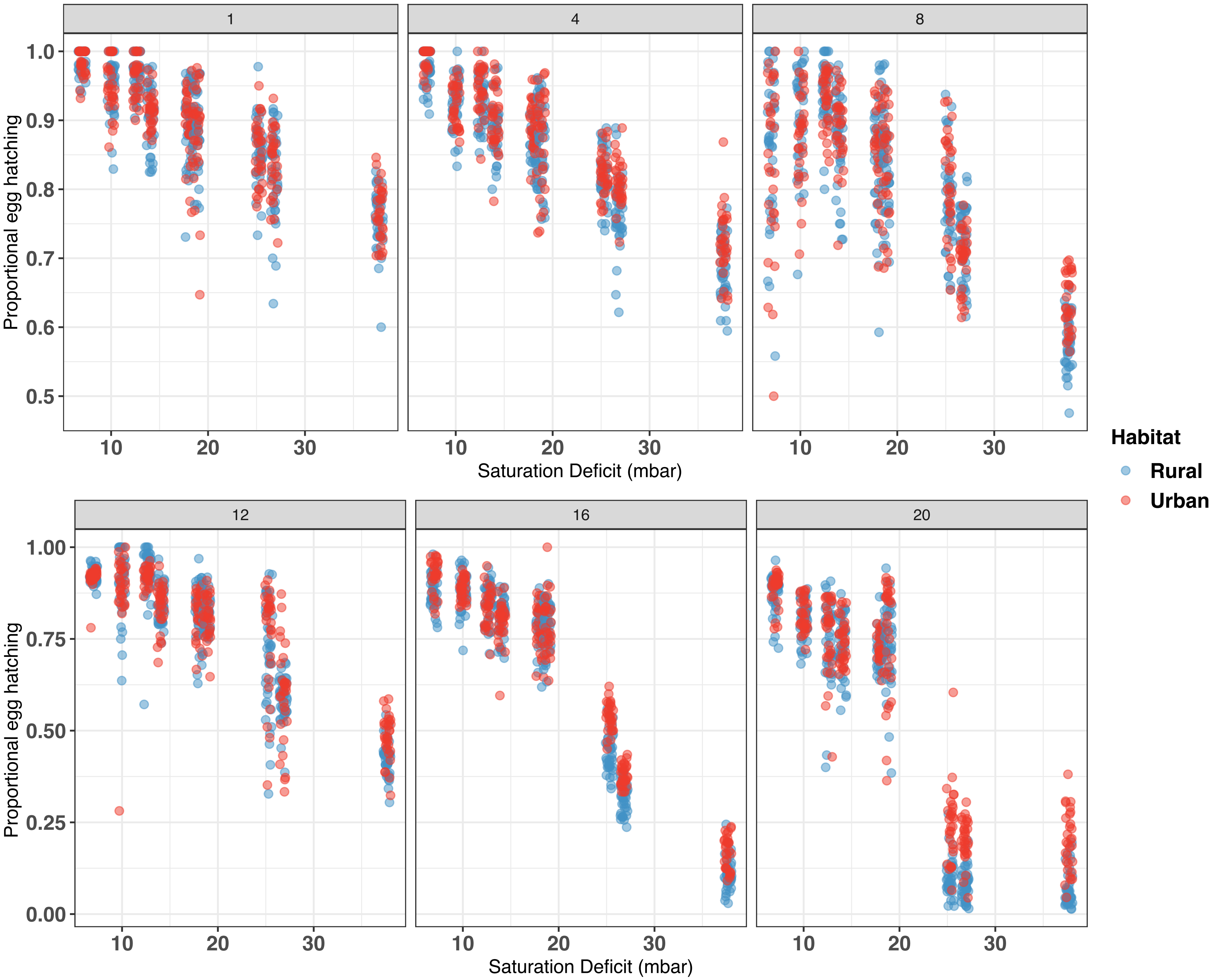

Supplementary Figure 5: Time to 50% and 75% egg viability loss across saturation deficit levels.

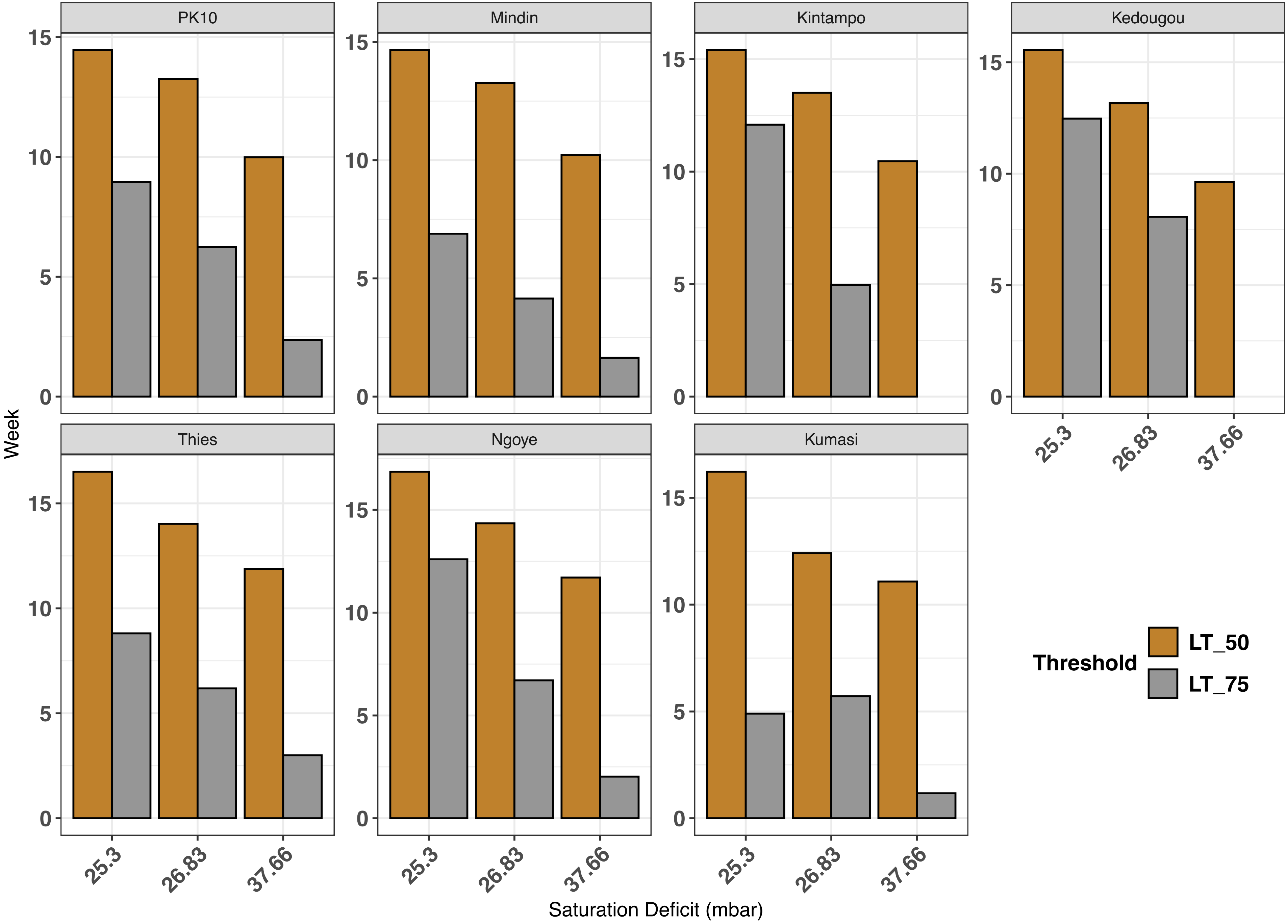

Supplementary Figure 6: Projected egg hatching probabilities under future climate scenarios across six African locations.

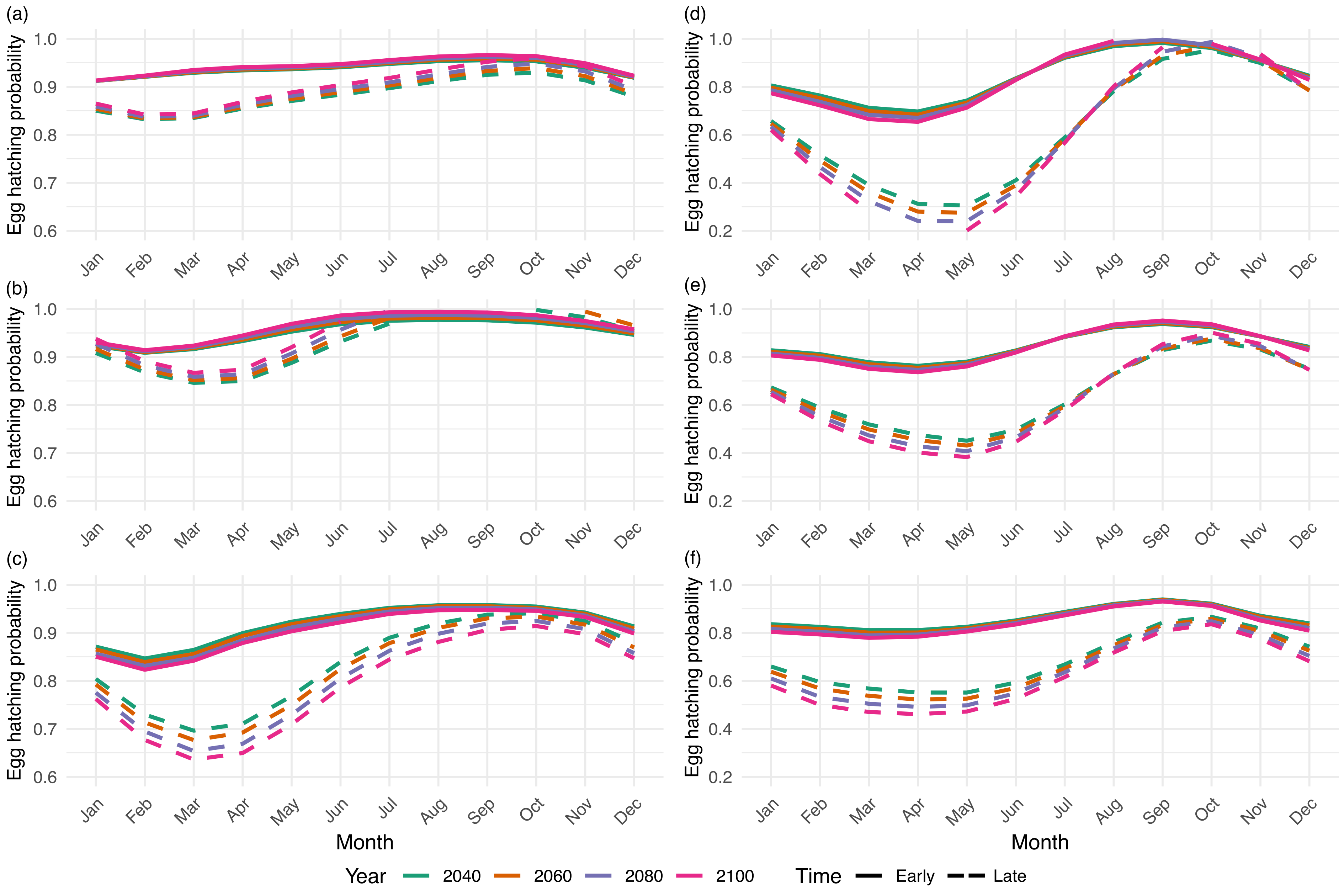

Supplementary Figure 7: Projected mosquito survival probabilities across six African locations under SSP3 climate scenario.

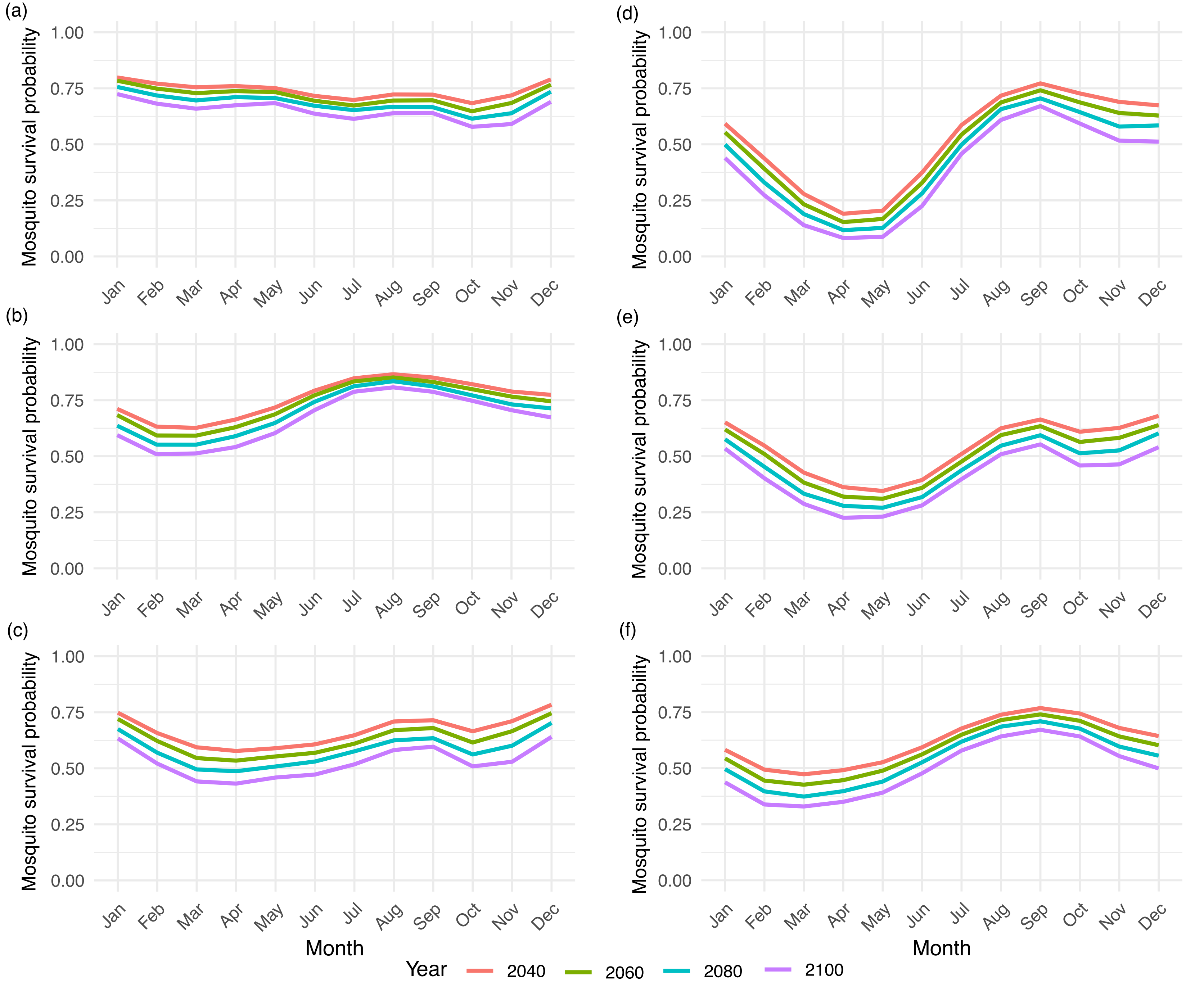

Supplementary Figure 8: Egg hatching probabilities across selected study locations under future climate projections.

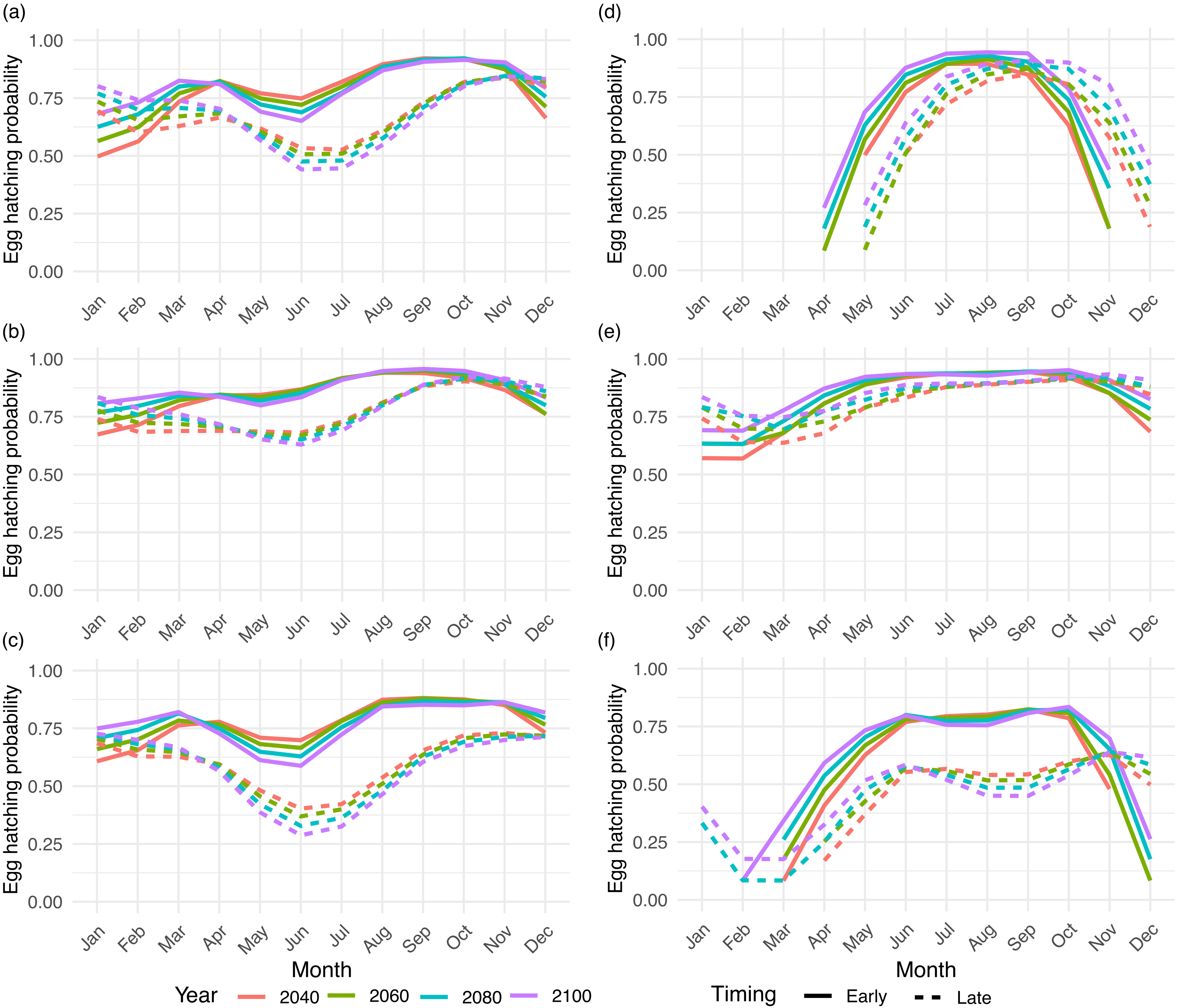
